# CCDC103-mediated assembly of the R2C complex links RUVBL1-RUVBL2 to Primary Ciliary Dyskinesia

**DOI:** 10.1101/2025.09.11.675549

**Authors:** Hugo Muñoz-Hernández, Maxwell Shelley, Skye Kelsall, Theodoros I. Roumeliotis, Jyoti Choudhary, S. Mark Roe, Laurence Pearl, Michal Wieczorek, Mohinder Pal

**Affiliations:** Department of Biology, Institute of Molecular Biology & Biophysics, ETH Zürich, Zurich, Switzerland; School of Natural Sciences, University of Kent, Canterbury, CT27NH, UK; Division of Structural Biology, Institute of Cancer Research, Chester Beatty Laboratories, 237 Fulham Road, SW3 6JB, London, UK; Division of Cell and Molecular Biology, Institute of Cancer Research, Chester Beatty Laboratories, 237 Fulham Road, SW3 6JB, London, UK; Genome Damage and Stability Centre, University of Sussex, Brighton, BN1 9RQ, UK; School of Life Sciences, University of Sussex, Brighton, BN1 9RQ, UK

**Keywords:** CCDC103, DNAAF19, RUVBL1, RUVBL2, Axonemal Dynein, HSP90, R2TP and RPAP3

## Abstract

Primary ciliary dyskinesia (PCD) is a genetic disorder caused by defective cilia motility, powered by axonemal dynein motors. Assembly of these motors is facilitated by the molecular chaperone HSP90, its co-chaperone RUVBL1-RUVBL2 and adaptor proteins such as CCDC103 (aka DNAAF19). Mutations in CCDC103 identified in PCD patients impair dynein assembly, contributing to the disease pathology. Here, we present the cryo-electron microscopy structure of the human RUVBL1-RUVBL2-CCDC103 complex at 3.2Å resolution, a chaperone assembly we refer to as R2C. It comprises a hetero-hexameric RUVBL1-RUVBL2 ring bound to three CCDC103 molecules via their RUVBL2-binding domains (RBDs), which have additional functions. Unlike RPAP3 of R2TP, a previously defined co-chaperone of HSP90, CCDC103 lacks a PIH1D1-binding motif and TPR domains, but its flexible N-terminal region regulates RUVBL1-RUVBL2 oligomerisation. Our characterisation of the R2C complex presented here enhances understanding of the intricate protein network involved in Hsp90-mediated assembly of dynein motors and how adaptor mutations contribute to PCD.

## Introduction

Cilia are hair-like projections on the surface of cells, and their dysfunction causes ciliopathies. Defective cilia affect multiple organs in the human body, including the lungs and heart^1,2^. Therefore, ciliopathies underlie several diseases, including primary ciliary dyskinesia (PCD) and congenital heart defects (CHD)^3^. PCD is an inherited, homozygous autosomal-recessive disorder that affects more than 1 in 15,000 births^4^. PCD patients manifest laterality defects and often more complex heterotaxy^4,5^.

Functional motile cilia are composed of a 9+2 microtubule arrangement, where nine microtubule doublets are organised in a circle around two microtubules at the centre of the cilium^6^ . These microtubules are associated with multi-subunit dynein motors^10^. These are large macromolecular protein complexes (>2MDa in size), comprising more than 20 different polypeptides^7^. These motors facilitate cilia motility and bending, which is essential for their physiological functions of clearing dust particles and aiding the movement of mucus^8^. Ciliary motility also plays a vital role in fertility^9^. Several research groups have elucidated the structures of assembled dynein motors and outlined their interactions with microtubules^10,11^. Recently, efforts have been made to investigate the dynein assembly pathway^12,13^. However, the process by which these motors are assembled in the cytoplasm and loaded onto microtubules remains poorly understood.

Undoubtedly, axonemal dynein motor assembly is a complex process, and several cell biology studies show the involvement of molecular chaperones, co-chaperones, and adaptor proteins, also known as dynein axonemal assembly factors (DNAAFs)^14–16^. Notably, it is known that either mutations in the genes encoding dynein proteins, or the chaperones/adaptors can disrupt cilia functions and lead to PCD ^4,16,17^. Among these chaperones and adaptors, Heat shock protein 90 (HSP90) and its co-chaperone R2TP (RUVBL1-RUVBL2-RPAP3-PIH1D1) have emerged as key players. These proteins form one of the most sophisticated chaperone machineries that recruit different adaptor proteins to assemble and activate a variety of client protein complexes, including dynein motors, PIKK kinases (mTOR, ATM, ATR, SMG1, and DNA-PKcs), and eukaryotic RNA polymerase II ^18–22^.

To understand the mechanism of R2TP, we previously defined this complex and its connection with HSP90^23,24^. In humans, R2TP is a ∼450kDa multi-subunit complex composed of four conserved proteins: RUVBL1 (RuvB like 1), RUVBL2 (RuvB like 2), RPAP3 (RNA polymerase II associated protein 3) and PIH1D1 (PIH1 domain containing protein 1) corresponding to <u>R</u>vb1p, <u>R</u>vb2p, <u>T</u>ah1p, and <u>P</u>ih1p in yeast^22^.

RUVBL1 and RUVBL2 are AAA-ATPases that are composed of three domains; domains I and III form the ATPase (Figure 1), which is regulated by domain II (also called DII)^25^. The ATPase domains of the RUVBL1 and RUVBL2 oligomerise to form alternating hetero-hexamers, with six ATP binding sites^26^. These hetero-hexameric rings of the RUVBL1-RUVBL2 complex can further oligomerise and form dodecamers by interacting via their six DII domains^26,27^. RPAP3, substantially larger than its yeast analogue Tah1p, consists of two TPR domains that bind the HSP90 dimer and a PIH1D1 binding motif, followed by a C-terminal RUVBL2-binding domain (RBD) ^23,28^. A schematic representation of these structural features is shown in Figure 1.

**Figure 1.**
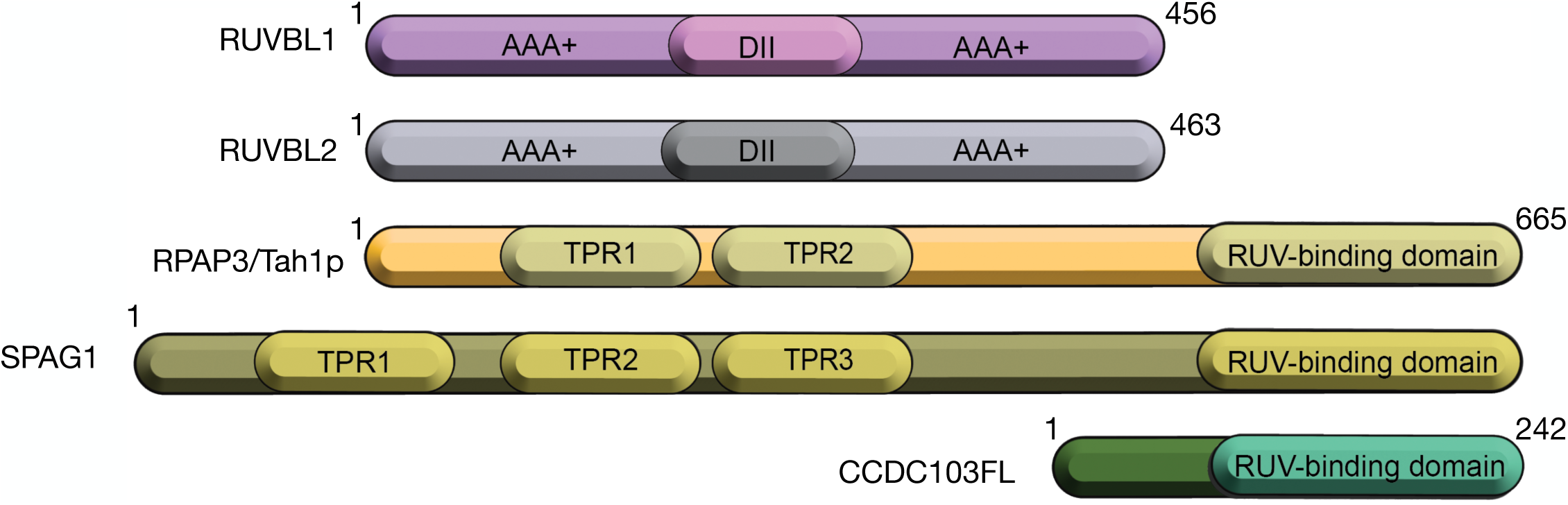
Schematic representation of the domain topology of the RUVBL1, RUVBL2, RPAP3, SPAG1 and CCDC103 proteins. RUVBL1 and RUVBL2 each contain AAA+ ATPase domain, which oligomerise to form a hetero-hexameric RUVBL1-RUVBL2 complex. The ATPase domain is separated by the DII domain, which is implicated in regulating the ATPase activity of the RUVBL1-RUVBL2 complex. RPAP3, a homologue of Tah1p, has two TPR domains and a C-terminal RUV-binding domain, also known as the RUVBL-binding domain, which is shared by other proteins, such as SPAG1 and CCDC103.

To assemble dynein motors, R2TP is implicated in binding to different adaptors such as coiled-coil domain-containing protein 103 (CCDC103, also known as DNAAF19) and SPAG1 (Sperm-associated antigen 1)^4,14,29,30^. Specifically, CCDC103 is required in the assembly of both inner dynein arms (IDAs) and outer axonemal dynein arms (ODAs) and has also been implicated in binding to microtubules in cilia^31^. Notably, CCDC103 and SPAG1 share similarities with the RPAP3 protein of R2TP, as both these adaptors contain the RBD domain first identified in RPAP3 (Figure 1). Clinically, mutations in CCDC103 provide strong evidence for its essential role in dynein assembly and pathogenic contribution in PCD. Several variants, pGly128fs25*, pHis154P and R35P have been reported in patients with infertility or *situs totalis* with supporting evidence from zebrafish studies^32^. Interestingly, the clinical manifestation of CCDC103 mutations can be variable, as recently reported in a case of the basketball athlete with a homozygous H154P mutation, who exhibited defects in axonemal dynein assembly in sperm without manifesting typical symptoms of PCD^33^. This variability could arise from tissue-specific expression of CCDC103, differences in its oligomeric states (monomer or dimer) and post-translational modifications^34^. Consistent with this, the H154P mutant CCDC103 has a variable effect on cilia motility compared to healthy individuals, as some patients with this mutation exhibit a normal cilium beating pattern within the same sample, while others display a reduced beating frequency^35^. Most studies indicate that mutations in CCDC103 destabilise the protein, leading to an impaired dynein assembly pathway, which contributes to PCD and infertility^36^. However, how CCDC103 is recruited to the R2TP and HSP90 system remains uncharacterised.

We have now demonstrated that CCDC103 directly binds to the RUVBL1-RUVBL2 complex, and we present here the first single-particle cryo-electron microscopy (cryo-EM) structure of the RUVBL1-RUVBL2-CCDC103 complex at a 3.2Å resolution. Therefore, we identified a variant of the R2TP-Like complex that we termed the R2C (RUVBL1-RUVBL2-CCDC103) complex. Similar to our previously determined cryo-EM structure of the R2TP complex, the CCDC103 protein possesses RBD domains that interact with the AAA-ATPase domain of RUVBL2 of the hetero-hexameric RUVBL1-RUVBL2 AAA-ATPase ring. Our high-resolution data allowed us to identify interacting residues in CCDC103 with RUVBL1-RUVBL2, which we mutated, with consequent disruption of the R2C complex. The N-terminal (1-99) residues of the CCDC103 appear to be flexible, and density for this region could not be located in the cryo-EM map. However, our cross-linking mass-spectrometry (XL-MS) data suggest that these N-terminal residues (1-99), which are rich in lysine, cross-link with the DII domains of the RUVBL1-RUVBL2. This data allows us to propose a model in which CCDC103 regulates RUVBL1-RUVBL2 oligomerisation. Taken together, our structural work, complemented by biochemistry and cross-linking mass spectrometry data, provides a platform for rational drug design to stabilise the CCDC103 variants and also to serve as a foundation for evaluating the pathogenicity of novel patient-derived variants in PCD patients.

## Results

### Cryo-EM structure of the R2C complex

To reconstitute the R2C complex, we first confirmed the direct interaction among its components using a pull-down experiment with a C-terminally strep-tagged RUVBL2 assembled with RUVBL1 (Figure 2A-B). We confirmed that the RUVBL1-RUVBL2-strep complex successfully co-precipitated the full-length CCDC103. Following these pairwise interactions experiments, we proceeded to reconstitute the R2C complex for cryo-EM analysis.

**Figure 2.**
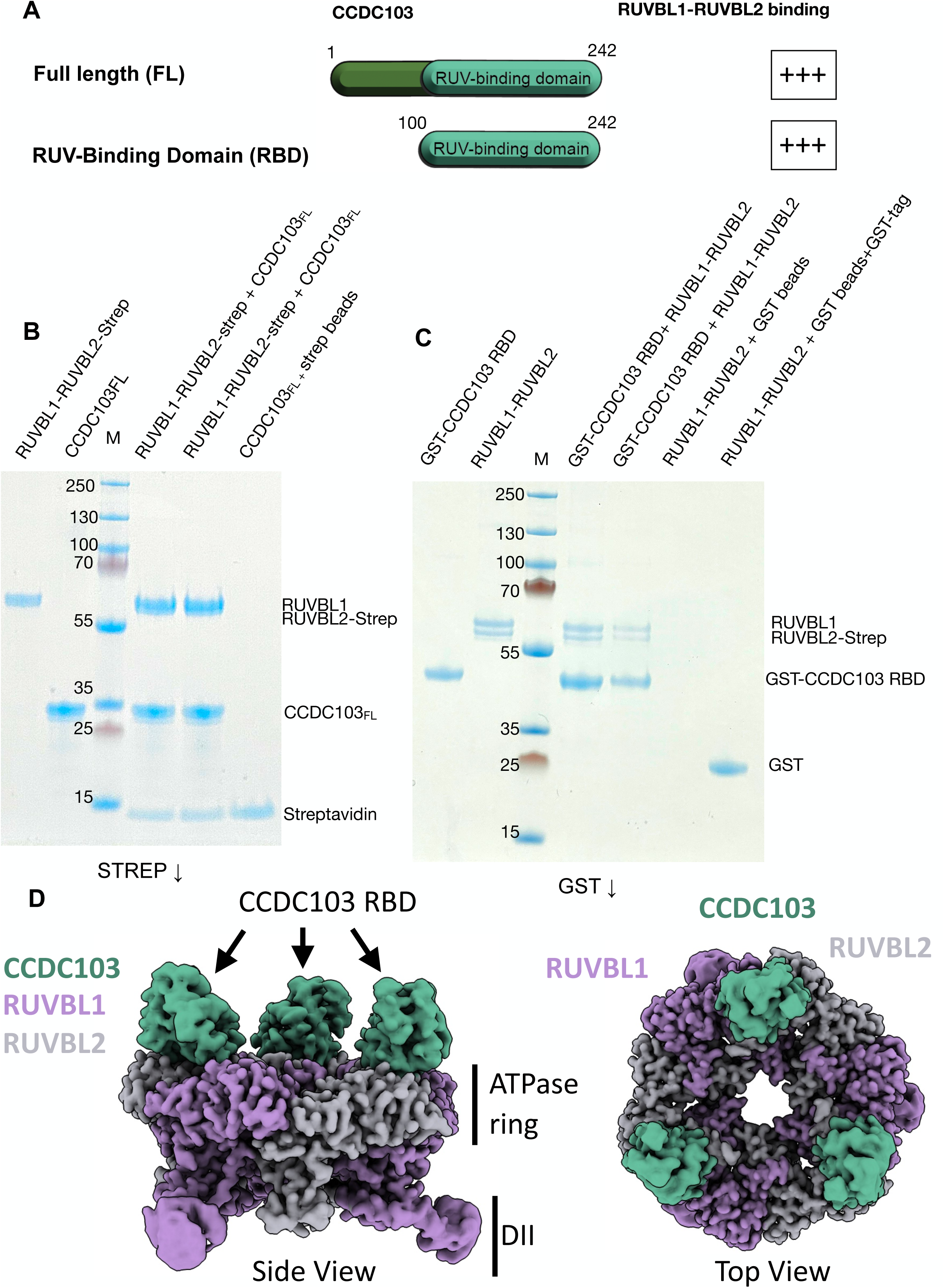
Pairwise interactions of CCDC103 with the RUVBL1-RUVBL2 complex and cryo-EM map analysis. A. Presents the outcome of the binding studies of CCDC103 full-length and the RBD domain of CCDC103 (RBD), which lacks the N-terminal (1-99) residues. Both constructs bind the RUVBL1-RUVBL2 complex, shown with positive signs. B. The pull-down experiments in which C-terminal strep-tagged RUVBL1-RUVBL2 complex successfully interacts with the full-length CCDC103. C. The GST-tagged RBD domain of CCDC103 is essential and necessary to bind the RUVBL1-RUVBL2 complex. D. The R2C cryo-map shows the presence of a hetero-hexameric RUVBL1-RUVBL2 ring decorated with the three RBD domains of the three CCDC103 molecules. The R2C complex displays both the side view and the top view, with RUVBL1-RUVBL2 forming the central part.

The R2C sample was vitrified by plunging into liquid ethane, and the cryo-data was collected with ThermoFisher Krios connected with the direct detector (K3 Gatan) (data collection parameters and model building are described in supplementary table 1). Initial 3D classification and refinement revealed that the density corresponding to the RBD domain of the CCDC103 was positioned on the ATPase ring of the RUVBL1-RUVBL2 complex, closely resembling our previously published cryo-EM structure of the human R2TP and R2TP-TTT complexes^23,37^. After further data processing, we observed a clear symmetric hetero-hexameric RUVBL1-RUVBL2 ring with its DII domains in an extended conformation, which is expected to hinder the formation of the dodecameric RUVBL1-RUVBL2 complex^23,25^. However, a small subset of 2D classes, corresponding to the dodecameric RUVBL1-RUVBL2 assembly in complex with CCDC103, was present, representing the assembly pathway of the mature R2C complex.

Subsequent 3D reconstruction of the data revealed the presence of the RBD domains of three CCDC103 molecules, each interacting with the three ATPase domains of RUVBL2 within the hetero-hexameric ring (Figure 2D). These findings confirmed that the ATPase ring of the RUVBL1-RUVBL2 complex was fully saturated with the CCDC103 molecules. Taken together, our data unambiguously established the stoichiometry of this R2C complex as 3:RUVBL1-3:RUVBL2-3:CCDC103. To obtain the high-resolution structure of the CCDC103 bound to the RUVBL1-RUVL2, we performed a two-step 3D classification and alignment of the RBD-RUVBL1-RUVBL2 ring, followed by symmetry expansion of the most homogeneous subset of particles (122,648 particles); a symmetry expansion strategy was similarly employed to determine a 1/3 fragment of the human R2TP^37^. This approach allowed us to reconstruct the complex at an overall resolution of 3.2Å. However, despite multiple attempts at focused and multibody refinement on the individual subregions of the cryo-map, no interpretable density was observed for the N-terminal 1-99 residues of CCDC103, which strongly suggests that this region is intrinsically flexible.

Based on these results, we hypothesised that the RDB domains of CCDC103 serve as the main docking modules for the recruitment of the RUVBL1-RUVBL2 ring. To test this hypothesis, we generated another construct in which the RBD of CCDC103 without the first 99 residues, was N-terminally tagged with glutathione S-transferase (GST). This GST-tagged RBD alone successfully co-precipitated the RUVBL1-RUVBL2 complex, indicating that the RBD domain is necessary and sufficient to pull down the RUVBL1-RUVBL2 complex (Figures 2A and C). This is fully consistent with our cryo-EM analysis. Notably, relative to the full-length CCDC103 protein, the isolated RBD domain significantly improved protein over-expression, yield and stability at room temperature and solubility at high concentration (20 mg/ml) (Supplementary Figure 1). It has been reported that the CCDC103 residues 94-242 form oligomeric species through self-interaction sites^38^. Consistent with this observation, we also observed a shoulder before the central elution peak of the CCDC103 RBD on size exclusion chromatography (Supplementary Figure 1), suggesting the presence of minor oligomeric species. However, no larger oligomeric assemblies were present in our protein purification steps and conditions and cryo-EM maps.

### Atomic detail shows residues of the RBD of CCDC103 engaged with RUVBL2

The high-resolution structure of the R2C complex was determined by collecting a total of 27,114 movies in two datasets using a ThermoFisher Krios G3i microscope connected to the direct detector (K3 GATAN). The data was processed and classified, resulting in a final high-resolution reconstruction of the complex at 3.2Å resolution with symmetry expanded particles (Supplementary Table 1, Figure 2 and Figure 3). We docked our previously published coordinates for the RUVBL1-RUBL2 complex^25,39^. For CCDC103, we utilised the model generated by the AlphaFold algorithm^40^ to guide the model building of the R2C complex assembly. The final refined structure of R2C revealed that RUVBL1-RUVBL2 forms the central part of the complex, revealing clear density for their ATPase domains and the protruding DII, particularly that of RUVBL1 (Figure 3A and B). However, the N-terminal region (1-99) of CCDC103 could not be located in the cryo-EM map. Notably, at the present resolution, atomic structural details of the ADP molecules and the side chain conformations of the residues of RUVBL1-RUVBL2 are well resolved in the cryo-EM map (Figure 3C and D).

**Figure 3.**
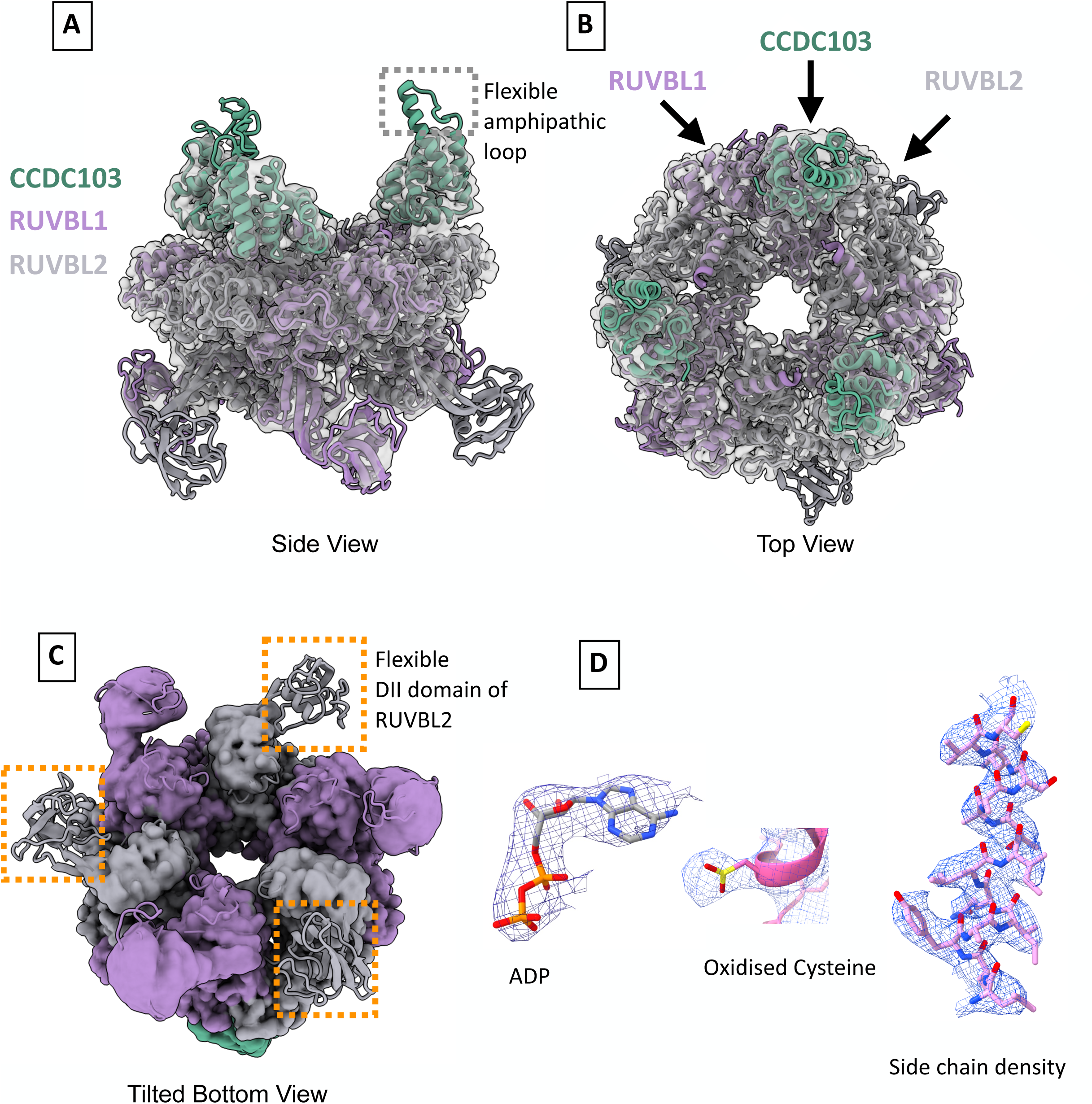
Cryo-EM structure of the R2C complex in detail. A. and B. show side and top views, respectively, of the R2C complex. This detailed structure shows that CCDC103 has a flexible amphipathic loop, which is absent in the RBD domain of RPAP3. C. The tilted bottom view of the R2C complex reveals that the binding of three CCDC103 molecules to the ATPase domains of the RUVBL2 components relays a conformational change, which renders their DII domains flexible. D. High-resolution features in the cryo-map, especially ADP molecules, oxidised cysteine 133 of CCDC103 and the side chains of the residues 399-413 of RUVBL2.

For RBD of CCDC103, we observed continuous density consistent with its predicted alpha-helical structure. However, an amphipathic loop in CCDC103, opposite to its RUVBL2 binding side, is highly flexible, and its corresponding density is poorly resolved (Figure 3A). As noted above, the R2C cryo-EM structure contains three CCDC103 molecules bound on top of the well-defined ATPase domains, which are formed by their DI and DIII domains of RUVBL2 in the hetero-hexameric ring. Strikingly, engagement of RBDs of CCDC103 appears to induce a conformational change in the RUVBL2 molecules that propagates from their ATPase domains to their DII domains (Figure 3C). This conformational change makes the DII domains of RUVBL2 flexible and unresolved in the map, while leaving the DII domains of RUVBL1 unaffected. The presence of the resolved density for an oxidised side chain of Cys133 in CCDC103 and the side chains of the residues of the protein further underscores the atomic detail of the R2C complex (Figure 3D and Supplementary Table 3).

Building on these high-resolution structural insights, we next compared the interactions of the RBD of CCDC103 with the RPAP3 of the R2TP complex with the RUVBL1–RUVBL2 ATPase ring. Previously, we and others ^23,41^ demonstrated that the residues of RBD of the RPAP3, especially methionine 626 (M626) and phenylalanine 630 (F630), form hydrophobic interactions with the valine 389 (V389), tyrosine 482 (Y482), and valine 423 (V423) residues of RUVBL2 within the hetero-hexameric ring (Figure 4A). To validate these interaction interfaces, we performed site-directed mutagenesis and generated M626A and F630R double point mutations in the GST-tagged RBD of RPAP3. While the wild-type GST-tagged RBD successfully co-purified RUVBL2, the double mutant failed to do so (Figure 4C).

**Figure 4.**
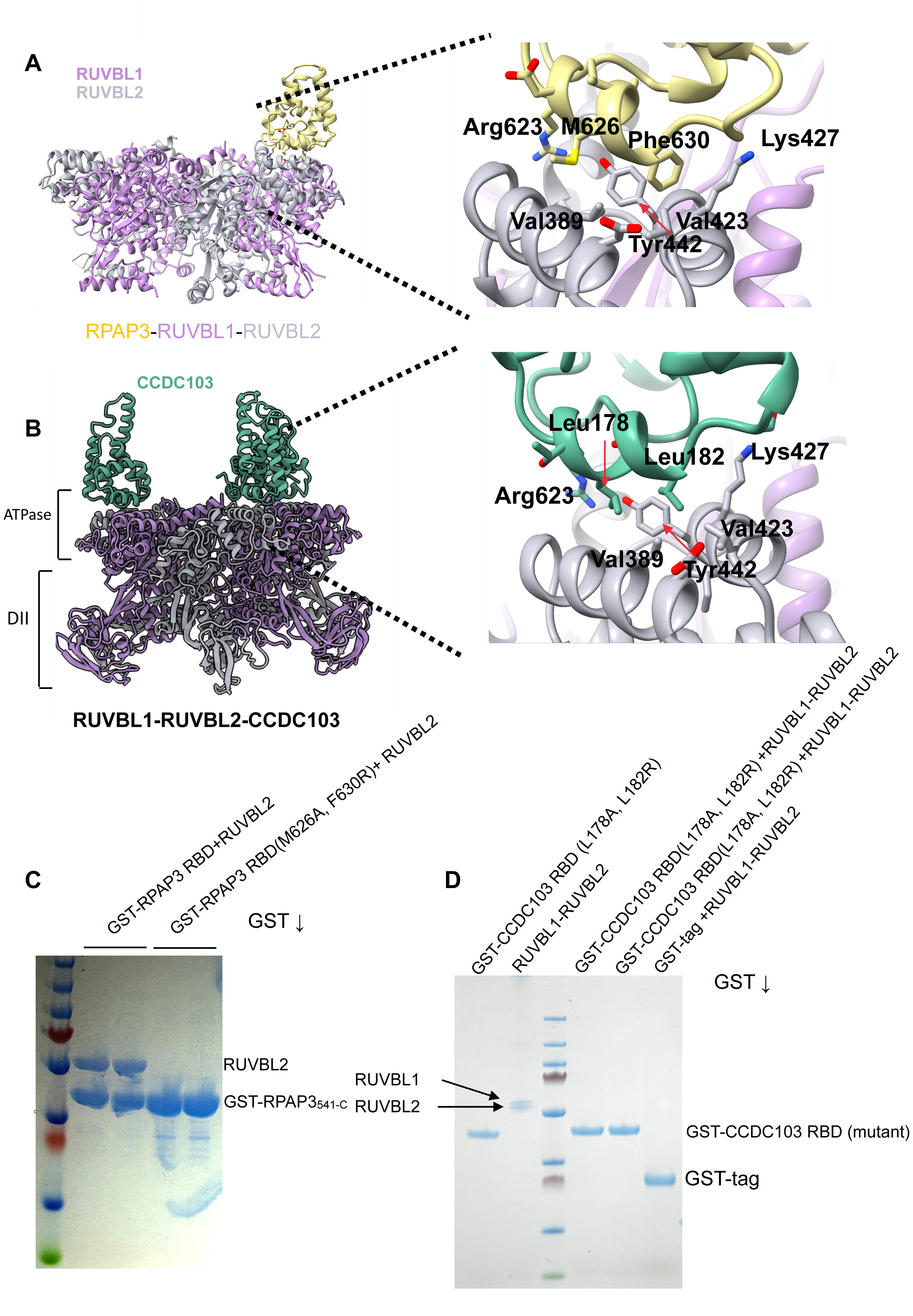
Atomic detail of the R2C structure and validation of the structure. A. The cryo-EM structure of R2TP (PDB code 6FO1), showing the interactions of the RBD domain of RPAP3 with the RUVBL2 in the RUVBL1-RUVBL2 complex. Specifically, Met626 and Phe630 of RPAP3 form hydrophobic interactions with Tyr442, Val389, and Val423 of RUVBL2. B. The cryo-EM structure of the R2C complex in which the RBD of CCDC103 engages RUVBL2 through interactions analogous to those formed by the RBD domain of RPAP3. Most of the interactions are hydrophobic in nature. In the RBD of CCDC103, M626 and Phe630 of RPAP3 are replaced with Leu178 and Leu182, respectively. C. and D. The pull-down experiments using GST-tagged RPAP3 and CCDC103 protein to validate the structure-based mutations that disrupt their interactions with RUVBL2. In C, the co-expressed wild-type GST-tagged RBD of RPAP3 pulled down RUVBL2, whereas the Met626Ala and Phe630Arg mutant protein was unable to do so. Similarly, the GST-tagged RBD of CCDC103 makes direct interactions with RUVBL1-RUVBL2 (Figure 2C), but the Leu178Ala and Leu182Arg mutant protein is unable to bind to the RUVBL1-RUVBL2 complex.

Similarly, in the presented R2C complex, the RBD of CCDC103 engages RUVBL2 through interactions analogous to those formed by RPAP3. In CCDC103, the equivalent hydrophobic residues are leucine 178 (L178) and leucine 182 (L182) (Figure 4 B). Clear density for these residues allowed us to interpret their interactions and guided us in generating structure-based mutations in the RBD of CCDC103 to disrupt binding to RUVBL2. We generated double point mutations (L178A and L182R) and tested their binding using GST-tagged pull-down assays. While the wild-type RBD of CCDC103 successfully pulled-down RUVBL2 (Figure 1C), the double mutant protein was unable to bind the hetero-hexameric ring (Figure 4D). Therefore, these interaction studies have defined the atomic interactions of CCDC103 with RUVBL2 within the RUVBL1-RUVBL2 complex.

### RBD of CCDC103 structurally diverges from RPAP3 and patient-derived mutations destabilise it

Although CCDC103 has a structurally homologous RBD domain to RPAP3, we discovered some notable differences between them. Specifically, CCDC103 has an additional amphipathic loop (Figure 5A), which is flexible in the cryo-EM map, and is entirely absent in RPAP3. This loop may represent a gain-of-function, enabling the R2C complex to either recruit another adaptor protein to facilitate axonemal dynein assembly or bind the dynein client directly. In contrast, the functions of RPAP3 could be as a general co-chaperone of HSP90 within the R2TP complex.

**Figure 5.**
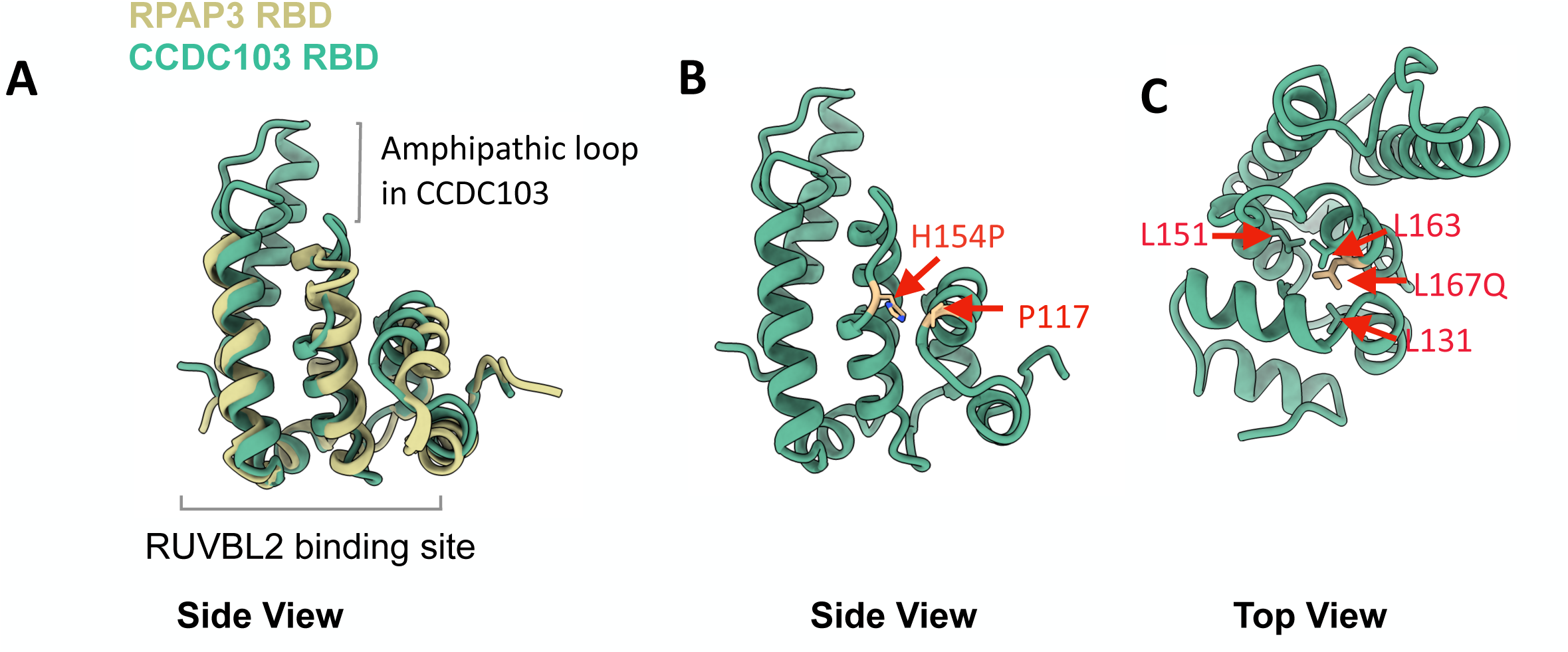
Structural comparison of CCDC103 with the RPAP3 and the location of patient-derived mutations in the RBD of CCDC103. A. A structural alignment of the RBD domains of the RPAP3 and CCDC103. Although the overall structural fold is very similar, CCDC103 has an additional amphipathic loop, which is flexible in the R2C complex but absent in RAPAP3. B. And C shows the location of patient-derived mutations in the RBD domain of CCDC103. In B. the His154, which is mutated to proline, is situated in the loop region and loosely packed against Pro117 of the adjacent helix. C. shows that Leu167, which is mutated to glutamine, is buried in a hydrophobic pocket formed by Leu131, Leu151 and Leu163. Our in-silico mutation in Leu167 to Gln creates steric clashes with the adjacent residues, therefore destabilising the CCDC103 protein.

The high-resolution structural detail also revealed that H154, which is mutated to proline in CCDC103 in some PCD patients, resides within the loop region, which was previously described to be in the alpha-helix^38^. Interestingly, this mutation is situated adjacent to this flexible amphipathic loop (Figure 5B), which is unique to CCDC103. Although our structural data lacks clear density for the sidechain of H154, our data suggests that the H154P mutation destabilises CCDC103. This is consistent with the previous observations that the proline insertion will alter backbone Φ angles (bond angles between the Cα-amide N bond) and Ψ angles (the Cα-carbonyl C bond), compromising protein stability^38^. Additionally, thermal shift assays demonstrate that the wild-type full-length CCDC103 and its isolated RBD domain are highly stable, with melting temperatures of 49 °C and 51°C, respectively (Supplementary Figure 5). In contrast, the H154P mutant protein is highly unstable and seems cytotoxic to bacterial cells, as it could not be successfully expressed and purified.

Since PCD and congenital heart defect (CHD) diseases are interconnected, we also identified a pathogenic and highly conserved L167Q variant of CCDC103 from the UK100K CHD patient cohort, which is also present in the genome aggregation database (gnomAD)^42^. Structural mapping of the L167 residue of CCDC103 in our recently determined cryo-EM structure shows that it is buried in a highly hydrophobic pocket formed by L163, L167, L151 and L131 (Figure 5C). Using in-silico substitution of L167Q using Pymol^43^ generates steric clashes with the surrounding residues, which strongly suggest that this mutation also destabilises CCDC103. These results further demonstrate that our structural data, complemented by biochemical analysis, provide a platform to predict the pathogenicity of genetic variants in both PCD and CHD patients.

### Flexible N-terminus of CCDC103 interacts with DII domains of RUVBL1-RUVBL2

Despite the high-resolution features present in the R2C complex, the highly flexible N-terminal residues (1-99) of CCDC103 could not be located in the cryo-EM map. To determine the spatial organisation of these residues in the R2C structure, we employed cross-linking mass-spectrometry (XL-MS), a strategy we previously utilised to complement previous cryo-EM work of the yeast and human R2TP complexes^23,24^. We utilised the reconstituted R2C complex for cross-linking mass spectrometry. This experiment yielded a total of 298 unique crosslinks, including 155 intra-protein and 143 inter-protein crosslinks (Figure 6A and Supplementary Table 2). A circular interaction map of CCDC103, RUVBL1, and RUVBL2 revealed enriched crosslinking between the N-terminal region of CCDC103 and the residues ranging from 200 to 300 of RUVBL2, which form its DII domains (Figure 6A). These domains are critical in regulating the ATPase activity as well as the dodecameric assembly of the RUVBL1-RUVBL2 complex. Similarly, dense crosslinking was observed between the N-terminal region of CCDC103 and residues 100-300 of RUVBL1, indicating multiple points of contact within the DII domains. Additionally, we detected increased crosslinking between the N-terminus of CCDC103 and the C-terminal regions of both RUVBL1 and RUVBL2, suggesting multiple contact points across distinct regions of the complex.

**Figure 6.**
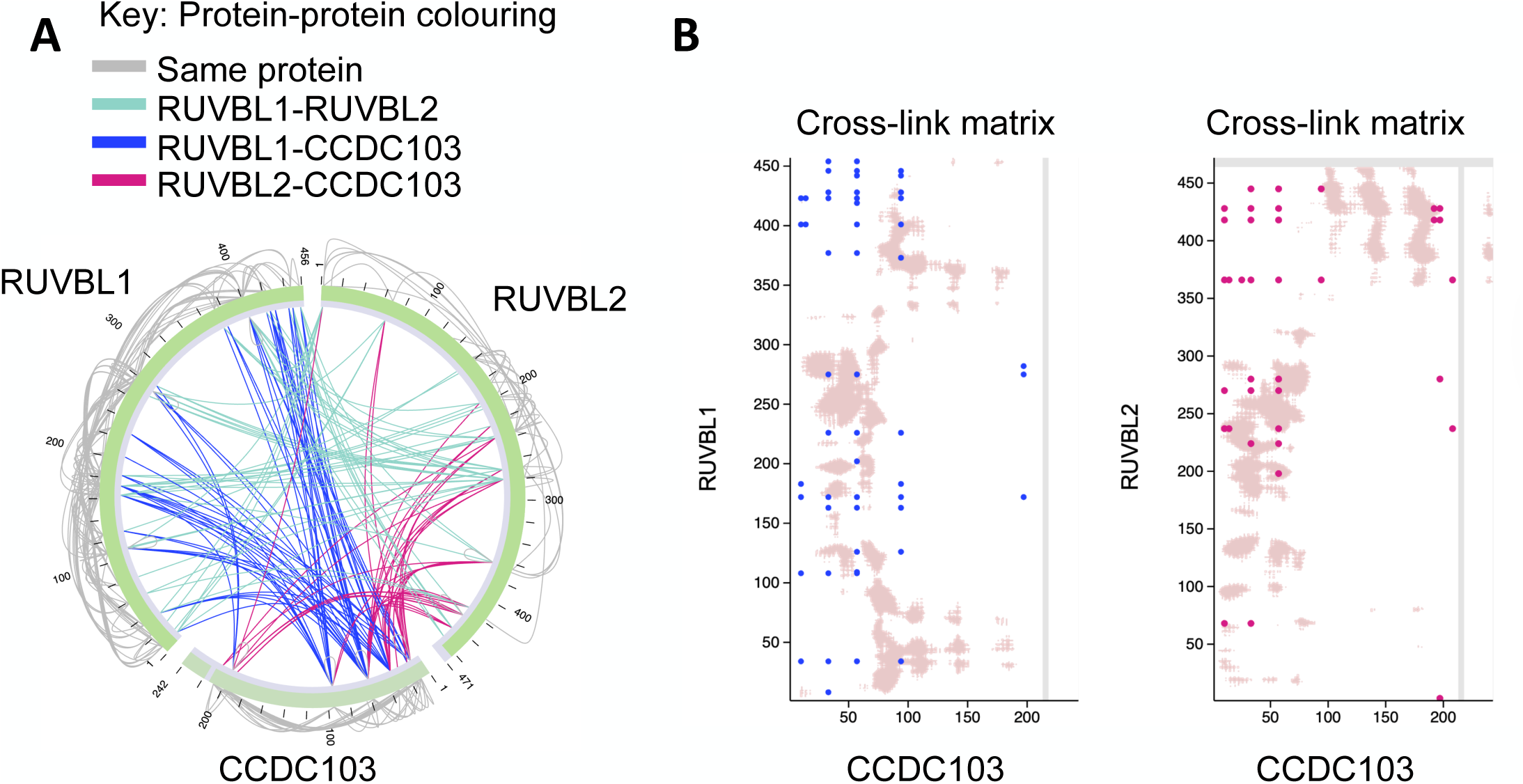
Cross-linking mass spectrometry of the R2C complex. A. A circular plot displaying all high-confidence crosslinks (FDR < 1%) involving CCDC103, RUVBL1, and RUVBL2, including both intra- and inter-protein crosslinks. B. 2D crosslinking matrices showing inter-protein crosslinks between CCDC103 and RUVBL1 (left panel) and CCDC103 and RUVBL2 (right panel). Crosslinks are mapped onto the linear sequences of each protein and highlight regions of increased crosslinking density, notably between the N-terminal α-helical region of CCDC103 and the 100-300 amino acid regions of RUVBL1 and RUVBL2. These are consistent with the spatial proximity seen in structural models shown in supplementary Figure 4A predicted using AlphaFold.

Mapping of the identified crosslinks onto the RUVBL1 and RUVBL2 (Figure 6B), followed by visualisation using 2D crosslinking matrices, revealed several crosslinks connecting the N-terminal region of CCDC103, containing a prominent α-helix, with the 200-300 amino acid region of RUVBL2. These crosslinks consistently overlapped with spatially proximal regions in the predicted AlphaFold model of the R2C complex (Supplementary Figure 4A-B), indicating a high degree of structural compatibility. This pattern strongly supports the presence of direct physical interactions between these domains and identifies them as likely contact sites within the complex. Similarly, crosslinks between the N-terminus of CCDC103 and the 100-300 amino acid region of RUVBL1 also showed satisfactory agreement with the structural predictions, further supporting the validity of the interaction and the conformational accessibility of the ATPase core domains in both RUVBL1 and RUVBL2 proteins.

Crosslinks involving the C-terminal regions of RUVBL1-RUVBL2 and the N-terminus of CCDC103 did not map within the predicted AlphaFold structure. This misalignment may be due to the low pLDDT scores shown in the AlphaFold models (Supplementary Figure 4A-B) for these C-terminal regions, indicating low-confidence structural predictions or intrinsic disorder. Such flexibility can permit transient or alternative conformations, which may bring these regions into contact and enable crosslink formation in solution, an event not captured in the single, static AlphaFold conformation. Together, our data indicate that the N-terminal region of CCDC103 directly contacts the DII domains of RUVBL1-RUVBL2. This interaction likely induces conformational changes in the DII domains, providing a structural basis for how CCDC103 binding disrupts the dodecameric assembly of the RUVBL1-RUVBL2 complex and promotes their hetero-hexameric state.

## Discussion

Primary ciliary dyskinesia (PCD) is a rare, genetically heterogeneous ciliopathy characterised by recurrent respiratory infections, laterality defects and infertility due to impaired motile cilia function^32,44^. PCD is frequently caused by defects in axonemal dynein assembly, which is a multistep process requiring the coordinated action of multiple assembly factors and chaperones^15,17,29,45,46^. Among these factors, CCDC103 has emerged as a key player as it has been implicated in dynein motor assemblies^31–33,35,38,47^. However, the mechanistic details of how CCDC103 participates in dynein arm assembly and how mutations in this factor lead to PCD pathology have remained poorly understood.

In this study, we present the first cryo-EM structure of CCDC103 in complex with the RUVBL1-RUVBL2 AAA+ ATPases, revealing the existence of a previously uncharacterized R2TP-like complex that we have termed the R2C complex. This discovery adds another critical layer to our understanding of the dynein assembly pathway. It also illustrates the versatility of RUVBL1-RUVBL2, which are at the centre of multi-protein chaperone and remodelling complexes. For instance, we previously showed that the RUVBL1-RUVBL2 complex directly binds TELO1-TTI1-TTI2 (TTT) adaptors, which specifically recruit PIKK kinases as client proteins^37^. In line with the literature and our previous discoveries, we now provide structural and biochemical evidence that CCDC103 directly interacts with RUVBL1-RUVBL2, which forms the core of the complex (Figure 2).

Our structural data advances research in the axonemal dynein assembly and PCD field by defining the oligomeric states of CCDC103 and the stoichiometry of the R2C complex. Previous studies have suggested that CCDC103 can form oligomeric states, which are essential for performing tissue-specific functions^34^. However, our structural data suggest that monomeric CCDC103 binds to RUVBL2 in the hetero-hexameric RUVBL1-RUVBL2 ring in a stoichiometry of 3RUVBL1:3RUVBL2:3CCDC103 (Figure 2D). This observation is consistent with our previous finding in the human R2TP complex, where the RBD domain in RPAP3 occupies all three positions on the RUVBL1-RUVBL2 ring^23,25^. Although the biological roles of the three copies of CCDC103 within the hetero-hexameric ring remain unclear, our findings contrast with the recent studies of the R2T complex from plants (*Arabidopsis thaliana)*^48^. In that system, the T component (atRPAP3), which contains the RBD domain, occupies only a single position per hetero-hexameric and dodecameric states of the atRUVBL1-atRUVBL2. This strict occupancy has been postulated due to allosteric communications between the RBD binding sites in the RUVBL1-RUVBL2 ring^48^, a phenomenon that is not observed in the R2C complex.

Although CCDC103 remains monomeric in our reported R2C complex, it plays a regulatory role in determining the oligomeric state of the RUVBL1-RUVBL2 complex. Previous studies have established that RUVBL1-RUVBL2 can exist in both hetero-hexameric and dodecameric states^48^; however, the precise role of these oligomers in the cell remains unclear. In the yeast R2TP system, Tah1p-Pih1p, analogues of RPAP3 and PIH1D1 in metazoan, facilitates the conversion of dodecameric Rvb1-Rvb2 to hetero-hexameric states by binding their DII domains^24,28,49^. Similar hetero-hexameric states are observed in the human R2TP complex, where RPAP3 positions PIH1D1 and its TPRs (Tetratricopeptide repeats) at the DII regions of the RUVBL1-RUVBL2^23^. Notably, although CCDC103 lacks the PIH1D1 binding motif, and the TPR domains typically associated with HSP90 recruitment, it nevertheless promotes a shift in RUVBL1-RUVBL2 equilibrium from a dodecameric to a hetero-hexameric state (Figure 3 A-C). This conclusion is supported by our cross-linking mass-spectrometry data, which show interactions between the N-terminal CCDC103 and the DII domains of the RUVBL1-RUVBL2 complex. Together, these findings provide a mechanistic insight into the CCDC103-mediated regulation of RUVBL1-RUVBL2 oligomeric transition (Figure 7).

**Figure 7.**
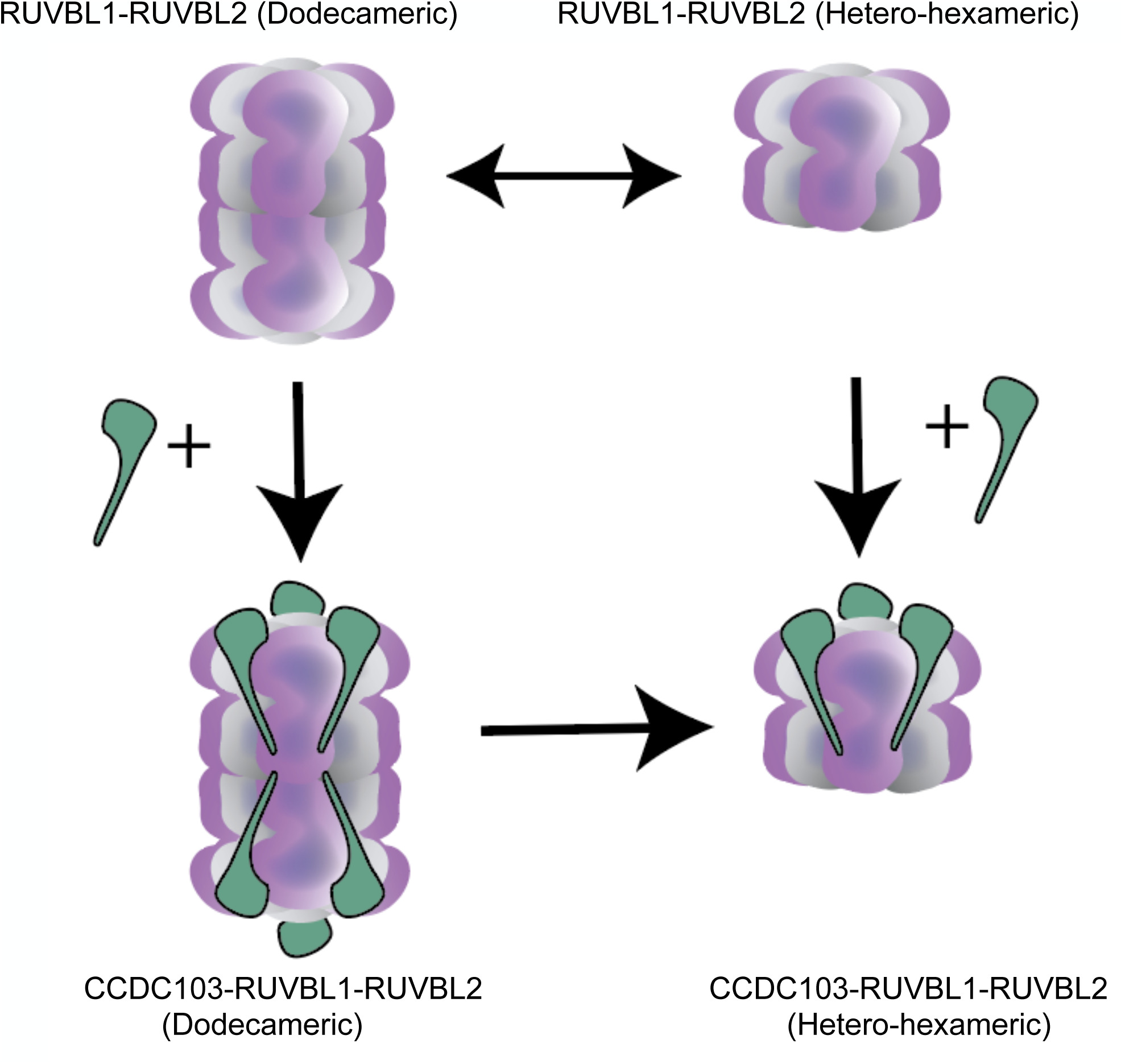
Proposed mechanistic insight of the R2C complex. The RUVBL1-RUVBL2 complex is known to transition from a dodecameric to a hetero-hexameric state. The addition of CCDC103 promotes shifts in equilibrium from the dodecameric to the hetero-hexameric R2C complex, as demonstrated by cryo-EM. This finding is supported by our XLMS data, in which the N-terminal flexible residues (1-99) of CCDC103 play a crucial role in shifting the dodecameric states of RUVBL1-RUVBL2 to the hetero-hexameric state by interacting with the DII domains of RUVBL1 and RUVBL2.

Based on the atomic resolution details in the R2C complex, we were able to selectively disrupt the interactions between CCDC103 and RUVBL1-RUVBL2 without compromising the structural integrity of the individual proteins (Supplementary Figure 1). However, the biological significance of disrupting these interactions remains to be evaluated, and future studies should be focused on determining how the loss of hetero-hexameric RUVBL1-RUVBL2 AAA ATPases affects dynein motor biogenesis. We have also mapped the patient-derived mutations on our recently determined R2C structure to assess their structural consequences. Unfortunately, H154P mutation lies within a flexible region. A possible explanation of this could be the dynamic nature of this region of the protein, or possibly another adaptor or client protein binding site, as H154 is situated next to the amphipathic loop, which is a unique feature of CCDC103, and is absent in the RBD of the RPAP3 protein. In contrast, the L167Q mutant appears to disrupt the overall fold of CCDC103. In summary, both mutations destabilise CCDC103 and contribute to its hypomorphic effect in PCD patients. Collectively, these insights underscore the potential of our structural work to provide a platform for translational research aimed at stabilising the mutant CCDC103 with small chemical ligands. Furthermore, this structural work may also serve as a predictive tool for evaluating the pathogenicity of novel patient-derived variants, thereby supporting more informed genetic diagnoses for patients with PCD.

Taken together, our results present the first cryo-EM structure of CCDC103 bound to the RUVBL1-RUVBL2 complex, forming the R2C complex. However, the mechanism by which R2C associates with the HSP90 molecular chaperone remains unclear, as CCDC103 lacks the tetratricopeptide repeat (TPR) domain typically required for HSP90 binding. This raises the intriguing possibility that an additional TPR-containing adaptor protein mediates the interaction between R2C and HSP90. Moreover, the identity of the axonemal dynein client proteins targeted by R2C remains unknown. Future studies are therefore essential to define these client interactions and to elucidate how R2C cooperates with other adaptors and assembly factors to coordinate the stepwise assembly of mature dynein motor particles.

## Materials and Methods

### Cloning

Flag-tagged-CCDC103, 6XHisRUVBL1 and RUVBL2-Step-tagged genes were cloned into pFBDM duet plasmids for protein expression in insect cells. These plasmids, containing the respective genes, were used to generate a bacmid in DH10bac cells. The white colonies containing the above genes were used for infecting the SF9 insect cell. The P1-P3 virus was propagated before large-scale protein expression. For pull-down, 6XHIS-RUVBL1-RUVBL2-strep genes were cloned into pRSETA and pCDF Duet 1 plasmids, respectively. The CCDC103 RBD domain was cloned into the pGEX6-P1 plasmid using *Bam*HI and *Eco*RI restriction sites, resulting in GST-tagged CCDC103 RBD.

### Bacterial Transformation

The 6XHIS-RUVBL1-RUVBL2-strep and GST-tagged CCDC103 RBD domain plasmids were used to transform *E. coli* Rosetta (DE3) pLysS competent cells (Novagen). Competent cells (25μl) were gently mixed with 2μl of plasmid DNA (100 ng/μl) in a sterilised microcentrifuge tube. The mixture was incubated on ice for 20mins followed by heat shock at 42°C for 60 seconds. The cells were immediately placed on ice for 2 minutes. 1 ml of sterile LB media was added to the cells, and the mixture was incubated at 37°C for 1 hour with shaking at 200 rpm. The 100μl of the transformed cells were spread onto LB agar plates containing 100μg/ml ampicillin and 100μg/ml spectinomycin. Plates were incubated at 37°C overnight, and the transformed cell colonies were obtained.

### Protein Expression

A single transformed colony was picked from an LB agar plate containing ampicillin and spectinomycin and inoculated into 50 ml falcon tube containing 10ml LB liquid media supplemented with 100 μg/ml ampicillin and 100 μg/ml spectinomycin. The culture was incubated overnight at 37°C with shaking at 200 rpm overnight. The overnight starter culture was used to inoculate 1L LB media containing respective antibiotics for protein expression in a 2L autoclaved conical flask. The culture was incubated at 37°C with shaking at 200 rpm until it reached an optical density at 600 nm of approximately 0.6, and the protein expression was induced by adding isopropyl β-D-1-thiogalactopyranoside (IPTG) to 1mM final concentration. The culture was then incubated overnight at 25°C with shaking at 200 rpm for high-level protein expression.

### Protein purification

The induced overnight bacterial cell mass was harvested by centrifugation at 7,459g for 20 minutes at 4°C using a JLA-9.1000 rotor (Beckman Coulter). For GST-CCDC103FL and GST-CCDC103 RBD, the cell pellet was collected and resuspended in 20 mM Tris-HCl, pH 7.85, and 500 mM NaCl. A protease inhibitor tablet (cOmplete, EDTA-free, Roche) was added, and approximately 25 ml of lysis buffer was used per litre of the original culture volume.

Cell lysis was performed by sonication using pulsed sonication consisting of 20sec on followed by 30sec off for 5 mins. The cell lysate was then centrifuged at 27,216g for 1hr at 4°C to pellet cell debris and insoluble material. The clarified supernatant containing the soluble protein fraction was collected for further purification. The GST-tagged CCDC103 protein was purified using glutathione Sepharose 4B resin (GE Healthcare). The bound protein was eluted with 50mM glutathione in 20 mM Tris-HCl, pH 7.85, and 500 mM NaCl. The purified protein was analysed with 4-12% SDS-PAGE, and the GST-tag was removed by adding 50μl(2mg/ml) of PreScion protease at 4 °C overnight. The GST tag was removed from the CCDC103FL and CCDC103 RBD proteins using a gel filtration column (HiLoad Superdex 75 16/60 column) equilibrated in 20mM Tris-HCl, pH 7.85, and 500mM NaCl buffer. The 6XHis-RUVBL1-RUVBL2 were purified using our previously defined protocol^23,25,37^.

### R2C protein complex reconstitution

The Bac-to-Bac system was used to generate the baculoviruses, which were used to infect Sf9 cells. The bacmid DNA samples were stored at −20°C. The propagated baculoviruses were stored at 4°C supplemented with 2% Foetal Calf Serum (Sigma-Aldrich cat # 12133C).

### *Spodoptera frugiperda 9 (Sf9)* cell growth conditions and protein purification

For the insect cell protein expression, Sf9 cells (*Spodoptera frugiperda 9*) were purchased from Thermo Fisher Scientific (cat# 11496015). Sf9 cell culture of 4 × 105 cells/ml was added to the 500ml of Insect-Xpress media containing Streptomycin/Penicillin (Lonza) antibiotics in a 2L roller bottle. The cell culture was incubated at 27°C with 150rpm shaking until the cell density reached 2 × 10^6^ cells/ml. Then the cells were then infected with titered viruses expressing proteins of interest. The infected cells were further grown for 72hrs at 27°C at 150rpm rotation. The cells were pelleted by spinning at 2,000*g* for 10min, at 4°C. The cell pellet was frozen at −80°C for long-term storage. For protein purification, 1L of cells expressing the R2C complex as well as Flag-tagged CCDC103FL were resuspended in 50mM Tris-HCl, pH 7.85, 300mM NaCl, 0.5mM TCEP. The cells were lysed using a homogeniser at 4°C. The cell lysate was clarified by centrifuging it at 17,000g for 1 hr. The supernatant was used to purify the R2C complex using a 5ml Streptactin column and the bound fraction was eluted using 5mM Desthiobiotin. For flag-tagged CCDC103, the protein was purified using anti-flag beads, and the bound protein was eluted with 0.5mg/ml 2X flag peptide.

### Pull-down experiments

10μM GST-CCDC103 RBD and 10μM RUVBL1-RUVBL2 were incubated with 70μl GST beads (GE Healthcare), which were equilibrated in 50mM HEPES pH 7.5, 140mM NaCl, 0.001% Tween 20 (pull-down buffer). For the negative controls for the RUVBL1-RUVBL2 complex, a 10μM of GST tag was used. The protein reaction mixture was incubated for 45 min at 4 °C, rotating at 20 rpm min^−1^. The beads were washed three times with 200μl of pull-down buffer, and the bound fractions were analysed by 4-12% SDS-PAGE (Invitrogen Ltd). Similar pull-downs were performed with full-length CCDC103 with RUVBL1-RUVBL2-strep using 50μl Streptavidin Mag Sepharose beads (Cytiva). The bound fractions were eluted using 2.5mM desthiobiotin in a pull-down buffer (Sigma-Merck Ltd), and the results were analysed by 4-12% SDS-PAGE (Invitrogen Ltd).

### Cross-linking mass spectrometry of the R2C complex

The protein complex was crosslinked with 1mM DSSO in a buffer containing 50mM HEPES, pH 7.85, and 140mM NaCl for 1hr, and the crosslinking reaction was quenched with 20mM ammonium bicarbonate. For trypsin digestion, triethylammonium bicarbonate buffer (TEAB) was added to the sample at a final concentration of 100mM. Proteins were reduced and alkylated with 5mM tris-2-carboxyethyl phosphine (TCEP) and 10 mM iodoacetamide (IAA) simultaneously and digested overnight with trypsin at final concentration 50ng/μl (Pierce). The sample was dried, and the peptides were fractionated with high-pH Reversed-Phase (RP) chromatography on a XBridge C18 column (1.0 × 100mm, 3.5μm, Waters) using an UltiMate 3000 HPLC system. Mobile phase A was 0.1% v/v ammonium hydroxide and mobile phase B was acetonitrile, 0.1% v/v ammonium hydroxide. The peptides were fractionated at 70μl/min with the following gradient: 5 minutes at 5% B, up to 15% B in 3 min, for 32 min gradient to 40% B, gradient to 90% B in 5 min, isocratic for 5 minutes and re-equilibration to 5% B. Fractions were collected every 100 sec, SpeedVac dried and pooled into 12 samples for MS analysis.

LC-MS analysis was performed on a Vanquish Neo UHPLC system coupled with the Orbitrap Ascend mass spectrometer (Thermo Fisher Scientific). Each peptide fraction was reconstituted in 30μl 0.1% TFA and 10μl was loaded to the PEPMAP 100 C18 5μm 0.3 × 5mm 1500 Bar trapping column. Peptides were then subjected to a gradient elution on a 25 cm capillary column (Waters, nanoE MZ PST BEH130 C18, 1.7μm, 75μm × 250mm) connected to a stainless-steel emitter on the Nanospray Flex ion source. Mobile phase A was 0.1% formic acid and mobile phase B was 80% acetonitrile, 0.1% formic acid. The separation method at flow rate 300nl/min was an 80 min gradient from 5%-35% B. Precursors between 380-1,400 m/z and charge states 3-8 were selected at 120,000 resolution in the top speed mode in 3 sec and were isolated for stepped HCD fragmentation (collision energies % = 21, 27, 34) with quadrupole isolation width 1.6 Th, Orbitrap detection with 30,000 resolution and 70ms Maximum Injection Time. Targeted MS precursors were dynamically excluded for further isolation and activation for 45 seconds with 10ppm mass tolerance. Identification of crosslinked peptides was performed in Proteome Discoverer 3.0 (Thermo Fisher Scientific) with the MS Annika search engine node for DSSO / +158.004 Da (K). Precursor and fragment mass tolerances were 10 ppm and 0.02 Da respectively with maximum 4 trypsin missed cleavages allowed.

Carbamidomethyl at C was selected as static modification and oxidation of M as dynamic modification. Spectra were searched against a FASTA file containing the sequences of the proteins in the complex concatenated with 1000 random UniProt E. Coli sequences as negative control. Crosslinked peptides were filtered at FDR<0.01 separately for intra/inter-links using a target-decoy database search. The crosslinked data was visualised using the xiVIEW platform^50^. Additionally, the Sequest HT node was used for identification of linear peptides. Dynamic modifications included Oxidation / +15.995 Da (C, M), Dioxidation / +31.990 Da (C), Trioxidation / +47.985 Da (C), DSSO Amidated / +175.030 Da (K), DSSO Hydrolysed / +176.014 Da (K) and Carbamidomethyl / +57.021 Da (C).

### Differential scanning calorimetry

1µl of the SYPRO orange dye (5000x, Sigma-Merck Ltd) was freshly diluted in 49µl of 50mM Tris-HCl, pH 7.85, 140mM NaCl to make 100x dye. 5µM CCDC103FL and CCDC103 RBD proteins were added to PCR tubes (Bio-Rad) and mixed with 1X Sypro Orange Dye. The reaction mixture (total 50µl) was incubated on ice for 10 mins. All the measurements were carried out in triplicate. The reaction mixtures were incubated in a qPCR thermocycler (Thermo Fisher QuantStudio™ 3 Real-Time PCR System). The protein melting curve was obtained by increasing the temperature from 5 to 95°C, as used in previously published research^51^. The final data was processed with Microsoft Excel.

### Cryo-EM sample preparation

EM grids were prepared with Quantifoil 300 mesh (copper, R 1.2/1.3), holy carbon grids. Grids were glow-discharged at 25 mA for 45 seconds. The RUVBL1-RUVBL2 and CCDC103 proteins were diluted in 25mM HEPES (pH 7.5), 140mM NaCl, and 10mM 2-mercaptoethanol–containing buffer containing 0.5mM ADP (pH 7.0) and incubated on ice for 180 minutes before Grid preparation. To prepare cryo-grids, 3.7μl of the protein complex was applied to the grid mounted on a Vitrobot Mark IV (ThermoFisher Scientific). The blotting of the grids was carried out for 3sec with blot force of -7 at 95% humidity and 6°C. Then, the grids were plunge-frozen in a liquid ethane/propane mixture and stored in liquid nitrogen until cryo-EM data collection.

### Cryo-EM data collection

A total of 27,114 movies were collected on two different grids using two TFS Titan Krios G3i at the ScopeM facility at ETH (Zürich, Switzerland 2). Both microscopes with FEG were operated with a Gatan K3 in CDS mode and a slit width of 20 eV on a GIF BioQuantum energy filter. These datasets were combined. Full cryo-EM data collection statistics and structure determination/refinement strategy are listed in Table 1 and Supplementary Figure 2. Automatic data collection was performed with “Faster acquisition mode” in EPU software (Thermo Fisher Scientific).

### Cryo-EM image processing

6,000 movies from each data collection were first processed to estimate a gain reference for each using “relion_estimate_gain”^52^. Movies were then imported into CryoSPARC for patch motion correction and patch contrast transfer function (CTF) estimation^53^. A blob picker with an elliptical blob size of 80-120Å was used to obtain a curated set of particles. The particles were selected based on 2D classification and visual inspection of micrographs, complemented with CTF fit resolution (2-18 Å), astigmatism (0-2000 Å), and relative ice thickness (0-1.8).

A ResNet16 neural network was trained on the curated particle set extracted from 16x downsampled micrographs with TOPAZ <u>(Bepler et al., 2020)</u>. TOPAZ models were generated for each independent dataset, and the software was used to pick and extract particles using a box size of 196. The TOPAZ-picked particles were subjected to 2D classification in CryoSPARC, and particles were selected based on the quality of 2D classes.

A total of 1,029,439 particles were selected from 2D classes and combined for five *ab initio* reference generation. One RUVBL1-RUVBL2 class was obtained, while the remaining four reference classes converged to noise. All particles were then subjected to 3D alignment by NU-refinement using the *ab initio* map as the starting reference. Subsequently, particles were re-extracted with a box size of 300 pixels and refined by applying local refinement focused on the AAA ATPase ring. A soft mask was used to encompass the RBD domains of CCDC103 on top of the RUVBL1-RUVBL2 ring.

A 3D classification was performed in CryoSPARC using a mask focused on the upper-ring density, including the RBD-like region. The classification was carried out into five classes with the resolution limited to 12Å, aiming to resolve the position and density of the RBD domain of CCDC103. The first (29%) and second (28%) classes both contained RBD-like density, but with a ∼60° mis-rotation relative to each other. The third class (26%) featured a RUVBL1_RUVBL2 class without the densities for the RBD domain of CCDC103. Particles from class 2 (291,028) were rotated and aligned to match class 1, after which the combined 587,687 particles from both classes were locally refined with Gaussian priors (10° SD on rotation and 4 Å SD on shifts), yielding a consensus and aligned R2C map.

The particles and the R2C consensus map were used for reference-based motion correction of both datasets, followed by particle re-extraction with dose-weighting applied. In total, 586,309 weighted particles were subjected to local refinement and CTF refinement. A subsequent round of 3D classification into five classes, limited to 8 Å resolution, was performed to assess heterogeneity and occupancy of the RBD domain density. A well-defined class 1 (23%; 139,290 particles) was then symmetry-expanded in C3 and locally refined, yielding a final R2C map at 3.2 Å resolution.

### Model building

For the R2C model, a structure prediction was first generated using AlphaFold 3 <u>(Abramson et al., 2024)</u>. The protein sequences of RUVBL1 (Uniprot Accession: Q9Y265), RUVBL2 (Uniprot Accession: Q9Y230), and CCDC103 (Uniprot Accession: Q8IW40) were used to assemble the initial model. The prediction was fitted into the consensus map and displayed in ChimeraX^54^ and then model built in Coot^55,56^. The resulting model was refined in real space using PHENIX^57^. The refinement statistics for models are available in Supplementary Table 1. Only RELION or CryoSPARC auto-sharpened EM density maps were used for final model refinements in PHENIX.

## Data availability

The EM maps have been deposited in the EM databank (http://www.emdatabank.org) with pdb code 9SN2 and map code EMD-55046.

## Acknowledgements

This work was supported by the Royal Society grant (RG\R2\232314), Action for A-T grant (24KEN01), UKRI BBSRC Impact Acceleration Awards (BB/X511158/1), UKRI MRC Impact Acceleration Award (MR/X502753/1) to MP, and the University of Kent start-up funds to MP. We are grateful for funding from the Wellcome Trust Investigator Award (210719/Z/18/Z/18/Z) to LHP in earlier stages of the work. This work was also supported by funding from ETH Zürich, an SNSF project grant (#310030_208120), and an SNSF starting grant (#TMSGI3_211309) awarded to MW. We also acknowledge access and support from the UKRI facilities at Diamond Light Source, Oxford.

## Author contributions

MP outlined the project and supervised the research. MP expressed and purified the R2C complex and performed the pull-down experiments to map interactions. MP, MS and SK designed the pull-down experiments. MP and HMH designed the cryo-EM experiments. HMH prepared the cryo-EM sample, collected cryo-EM data and determined the high-resolution cryo-EM map of the R2C complex. HMH refined and deposited the R2C model and cryo-EM maps. MP generated the structure-based site-directed mutagenesis in CCDC103, purified the mutant CCDC103, conducted pull-down experiments and carried out thermal shift assays. MP and MS processed the DFS data, and SMR helped in model building. TR and JC performed XL-MS, and LP provided intellectual input, data processing and data interpretation support. MP and HMH helped analyse and interpret results, analysed the data, wrote the manuscript and prepared the figures. LP and MW provided an overview of the project and their insight into the project. Everyone helped with manuscript correction and provided their insight.

## Key resources table

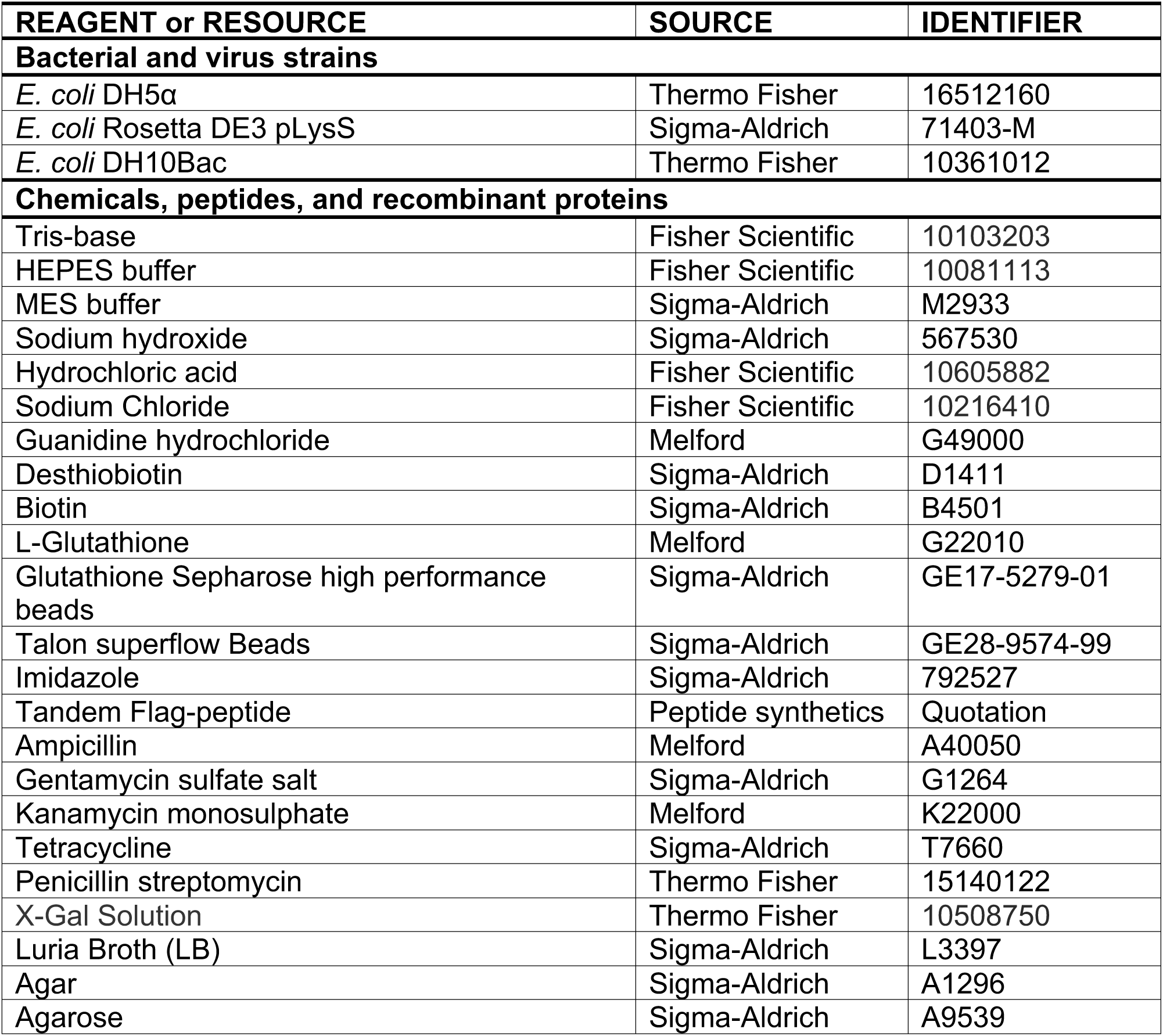
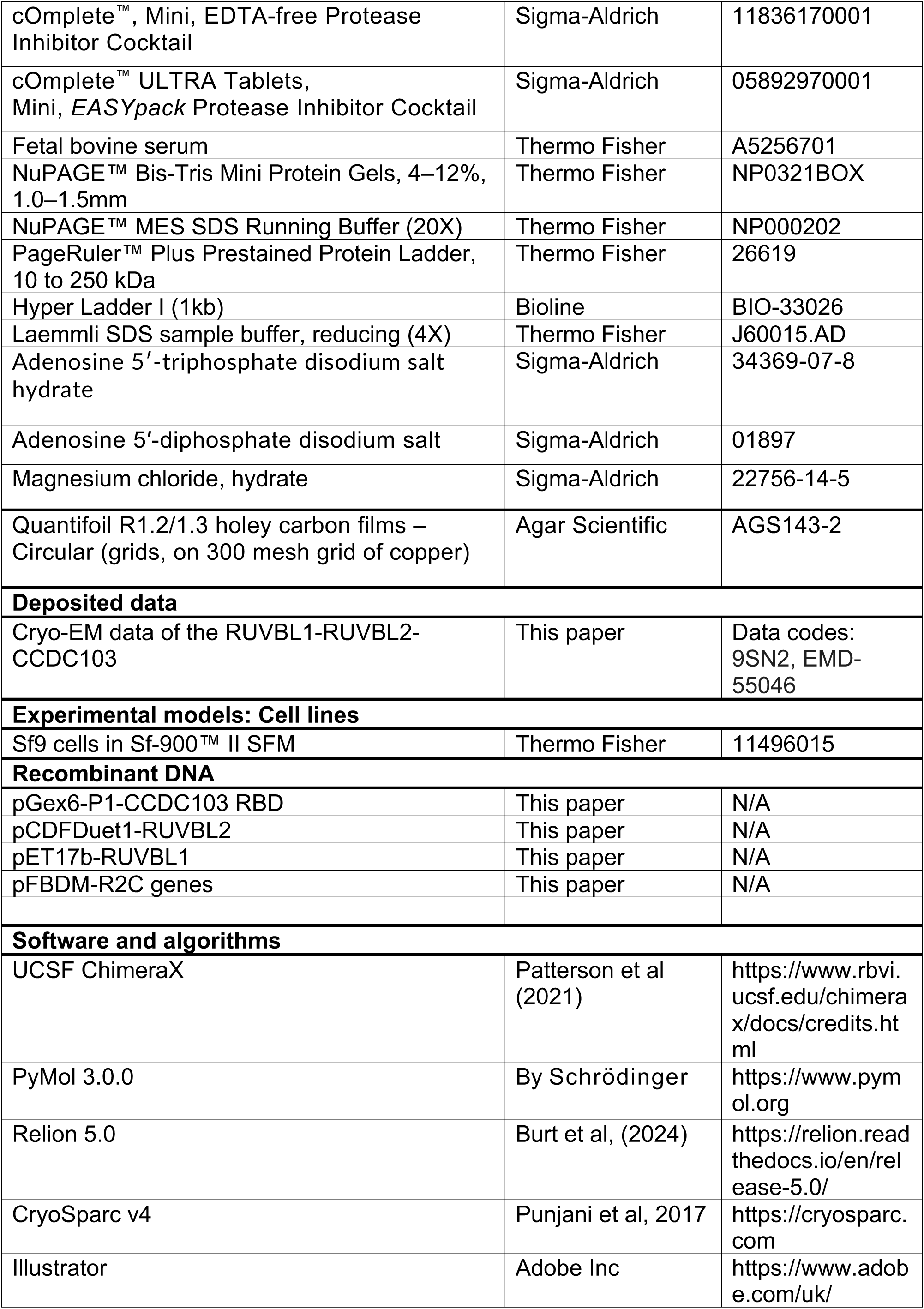
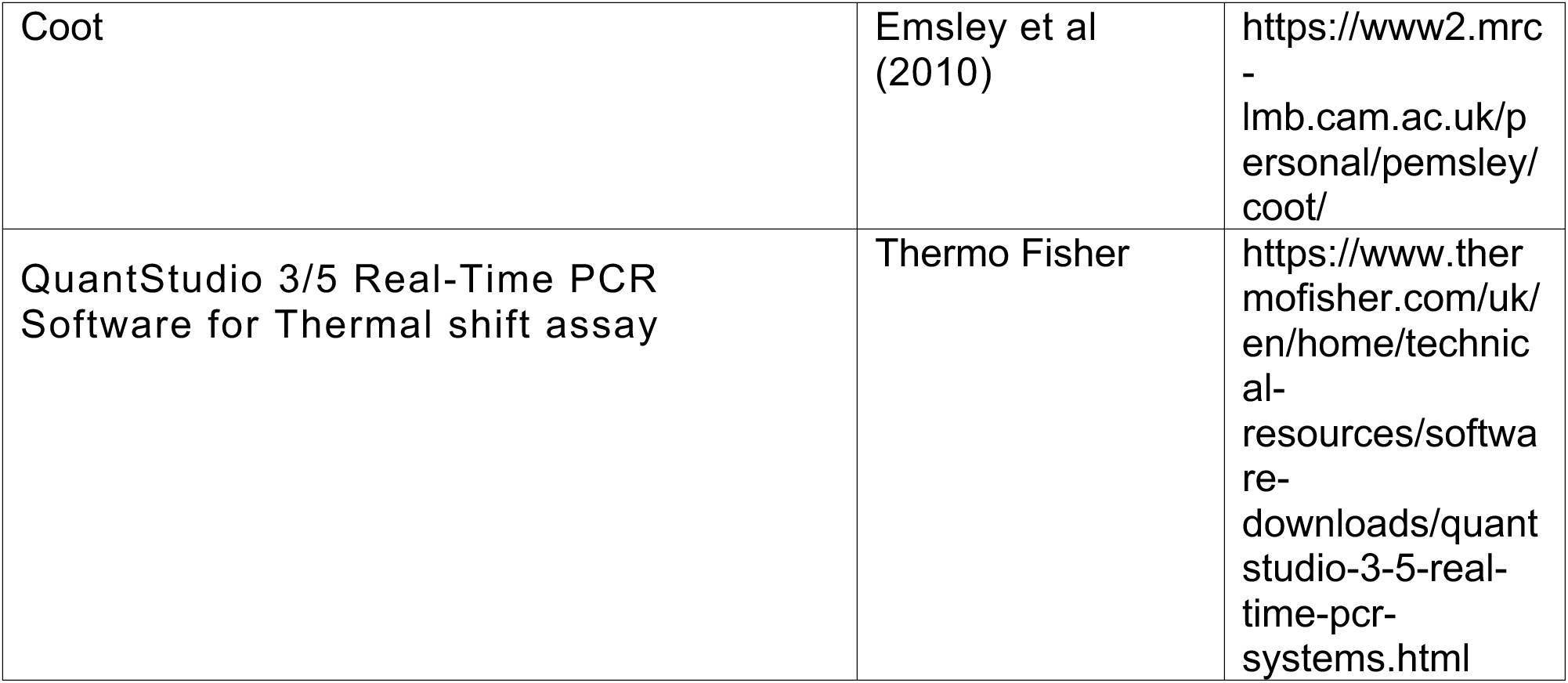

## Figure Legends

**Supplementary Figure 1. (S1).**
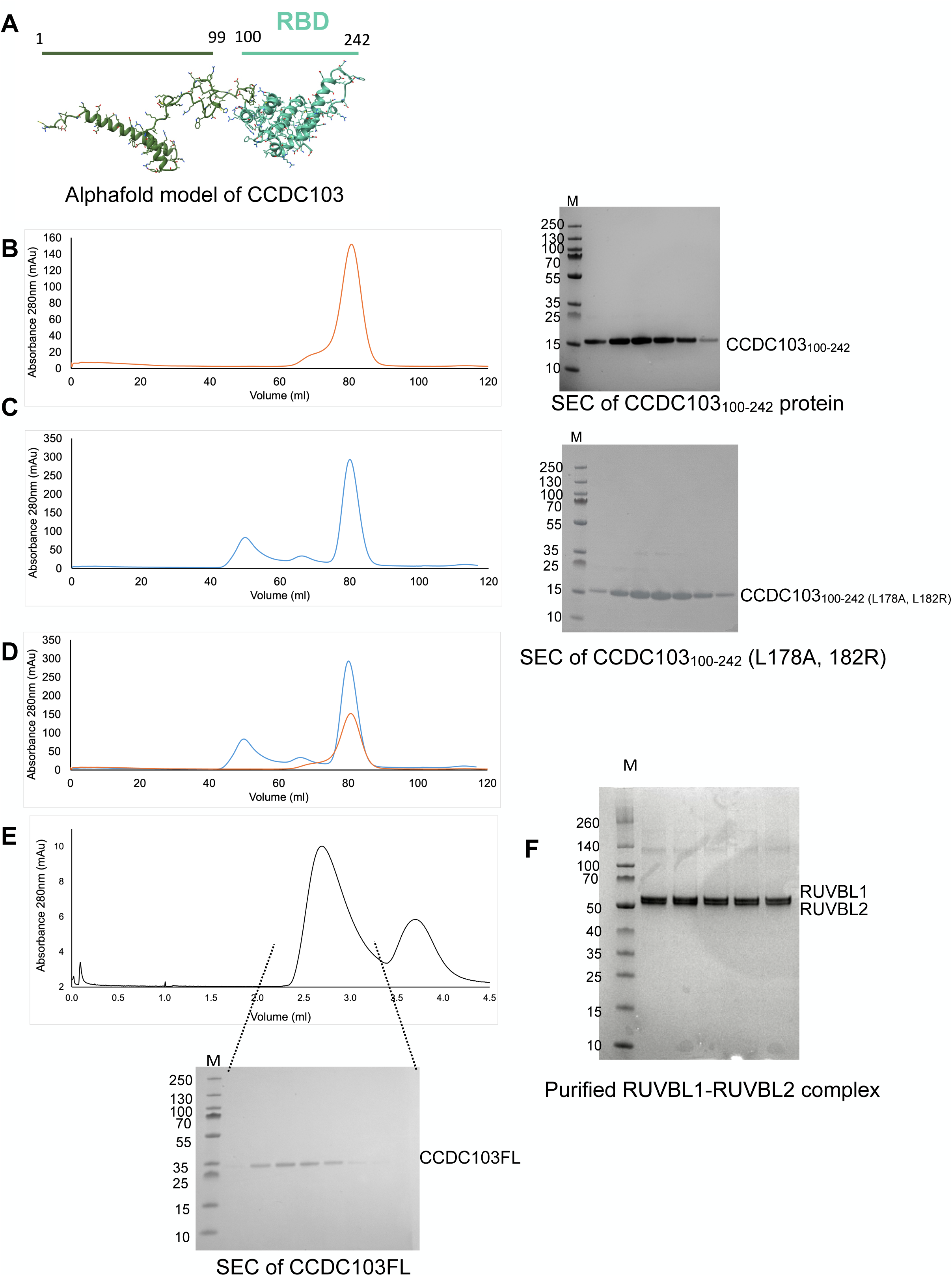
Protein purification. A. Domain mapping of RBD in the CCDC103 using the AlphaFold server. B. C. and D. depict the size exclusion chromatograms demonstrating that the wild-type and mutant RBD domains of CCDC103 have an equivalent profile as the protein elutes in a single peak with an elution volume of around 80ml using a Superdex 200 16/60 column. This is consistent with the structure-based mutations in RBD (L178A and L182R), which did not significantly alter the protein structure. The purity of these proteins was assessed by using a 4-12% PAGE. C. SEC purification of full-length CCDC103 protein on a Superose6 5/150 column. The quality of the protein was assessed by 4-12% SDS-PAGE. D. The RUVBL1-RUVBL2 complex was purified as we described in our previous publication of the cryo-EM structure of the human R2TP complex^23,25,37^. The purity of the protein complex was assessed using 4-12% SDS-PAGE.

**Supplementary Table 1. (ST1).** Cryo-EM data collection parameters and model refinement.

| **Category** | **Details** |
| --- | --- |
| Sample | RUVBL1-RUVBL2-CCDC103 (R2C) complex |
| Microscope | ThermoFisher Krios |
| Detector | K3 GATAN |
| Voltage (kV) | 300 |
| Nominal magnification | 81,000 |
| Electron exposure (e<sup>-</sup>/Å<sup>2</sup>) | 55 (dataset 1) and 48 (dataset 2) |
| Pixel size (Å/px) | 1.06 |
| Defocus range (μm) | -0.8 -2.2 |
| Initial particle images (no.) | 1,029,439 |
| Final particle images (no.) | 139,290 |
| Symmetry imposed | C1 with symmetry-expanded particles (120° rotation: 417,870 particles) |
| Map resolution (Å) | 3.2 |
| FSC threshold | 0.143 |
| Map resolution range (Å) | 2-8 |
| Map sharpening B factor (Å<sup>2</sup>) | -131.4 |
| Initial model used | AlphaFold 3 – 3315 Residues |
| N° of chains | 9 |
| Protein residues | 2526 |
| Ligands | ADP:6 |
| Bond lengths (Å) | 0.004 |
| Bond angles (°) | 0.847 |
| MolProbity Clashscore score | NONE |
| Ramachandran favored (%) | 95.17 |
| Ramachandran allowed (%) | 4.59 |
| Ramachandran outliers (%) | 0.24 |
| Rotamer outliers (%) | 1.76 |
| CC mask | 0.7951 |
| CC volume | 0.7734 |
| EMDB code | EMD-55046 |
| Accession number | PDB 9SN2 |

**Supplementary Figure 2 (S2):**
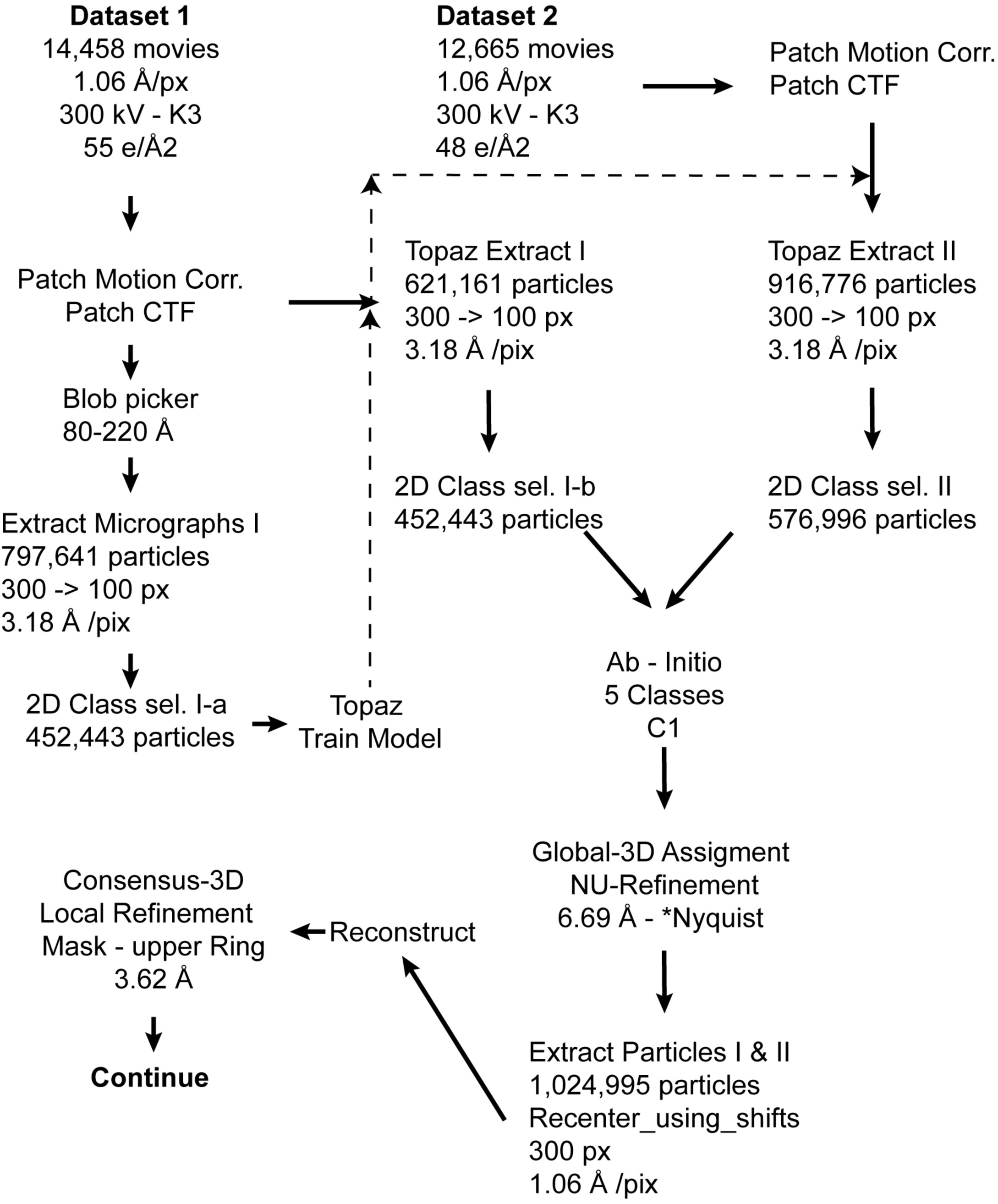
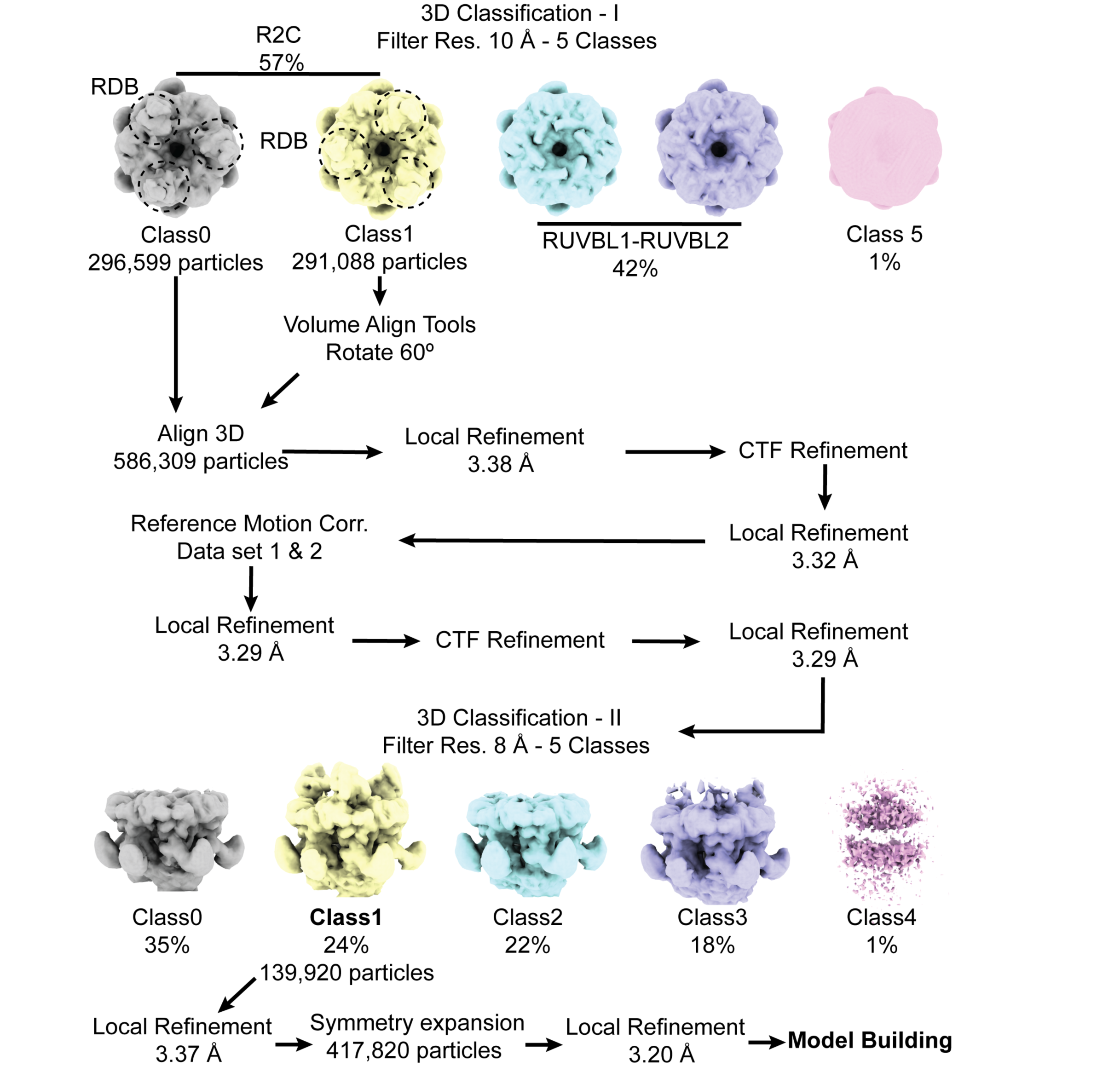
Cryo-EM data processing strategy to determine the high-resolution structure of the R2C complex.

**Supplementary Figure 3 (S3).**
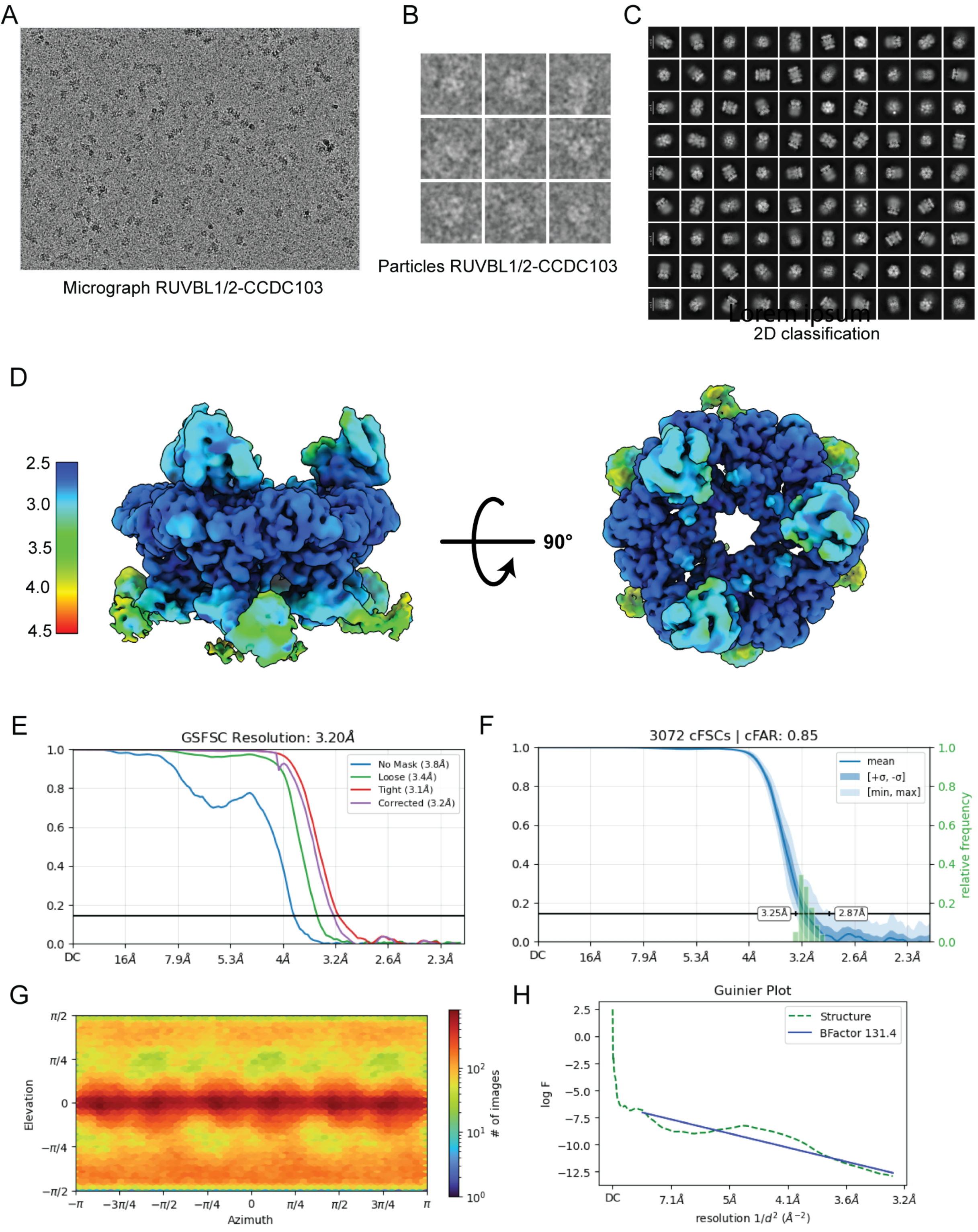
Cryo-EM data processing strategy to determine the high-resolution structure of the R2C complex. A-B shows the micrographs with protein particles, C. 2D classification showing the different views of RUVBL1-RUVBL2 and R2C complex, D. show the side and top views of the R2C complex. E. and F. show the FSC curve for resolution calculation. G and H. The particle distribution and Guinier plot.

**Supplementary Figure 4. (S4).**
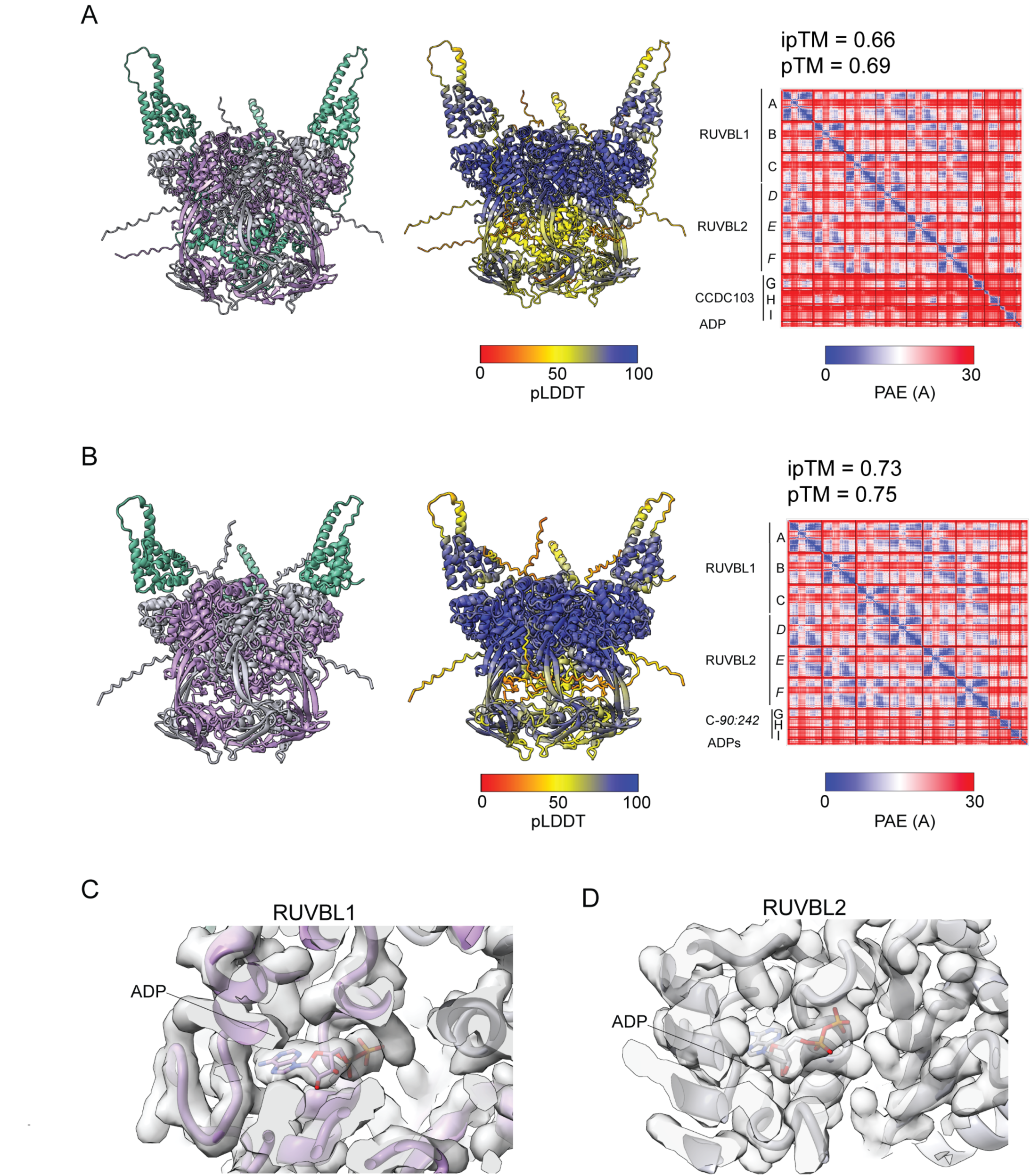
AlphaFold prediction and the cryo-EM map: A. and B. show the AlphaFold prediction of the R2C complex and partial alignment error plots. Although the R2C structure was very similar to the predicted AlphaFold models, the conformation of the DII domains of the RUVBL1-RUVBL2 is conformationally different from the cryo-EM map. C. and D. show that ADP molecules are loaded in the RUVBL1 and RUVBL2 molecules of the R2C complex, determined using the cryo-EM technique.

**Supplementary Figure 5 (S5).**
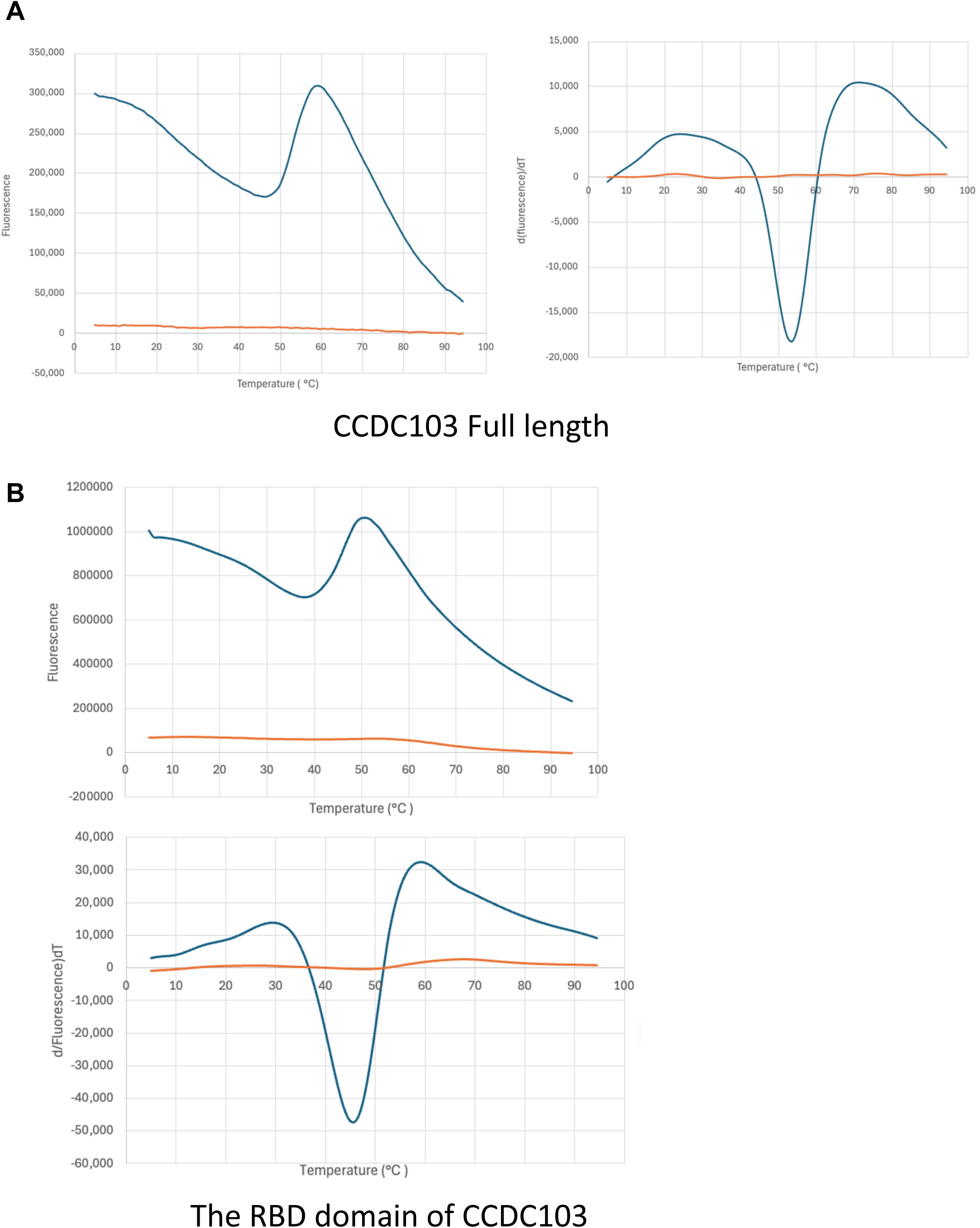
Differential Scanning Fluorimetry (DSF) of CCDC103 to determine melting temperature. A. The melting curve of the CCDC103FL protein, Tm ∼ 51°C. B. The melting curve of the RBD domain of CCDC103, Tm ∼ 49°C.

**Supplementary Table 2. Crosslinked peptides of the R2C complex identified by cross-linking mass spectrometry.**

**Supplementary Table 3. Identification of regular peptides in the R2C complex**

